# Mismatch-Assisted Toehold Exchange Cascades for Magnetic Nanoparticle-Based Nucleic Acid Diagnostics

**DOI:** 10.1101/2025.03.14.643268

**Authors:** Rebecca Sack, Joshua Evans, Florian Wolgast, Meinhard Schilling, Thilo Viereck, Petr Šulc, Aidin Lak

## Abstract

Sensitive, simple, and rapid detection of nucleic acid sequences at point-of-care (POC) settings is still an unmet quest. Magnetic readout assays combined with toehold-mediated strand displacement (TMSD)-based circuits are amplification- and wash-free, essential features for contributing to this demand. Nevertheless, nonenzymatic TMSD circuits are slow with low sensitivity for early disease diagnostics. Here, we propose novel mismatch-assisted toehold exchange (MATE) magnetic cascades, wherein magnetic susceptibility increases by dissociation of magnetic nanoparticles (MNPs) from engineered magnetic clusters upon detecting nucleic acid target in solution. The MATE relies on the generation of an allosteric toehold (TH) by spontaneous dissociation to efficiently recycle the target, amplify magnetic signal output, and enhance the assay’s kinetics. We show that introducing a mismatch in the allosteric TH domain enhances the overall declustering kinetics 7-fold, as also confirmed with oxDNA simulations, with the largest effect gained for the mismatch closest to where the branch migration by the target ends. By integrating MATE into magnetic diagnostics cascades, we demonstrate 3.6-fold better limit-of-detection (LoD) and 12-fold shorter assay time compared to our previous circuit design. Our work makes a major leap towards bringing MNP-based diagnostics much closer to the clinical POC settings by offering a simple, rapid, isothermal, and nonenzymatic assay workflow.

## INTRODUCTION

Certain nucleic acid sequences are highly specific to bacterial^1,2^ and viral^3–6^ genomes as well as cancer,^7–10^ and are thus the main target for early and accurate disease diagnostics. Reverse transcription polymerase chain reaction (RT-PCR), the ‘gold standard’ of molecular assays, can detect few nucleic acid copies through enzymatic amplification reactions.^11,12^ However, PCR requires expensive equipment and reagents and is timely and prone to contamination, making its application in point-of-care (POC) settings impossible. Particularly during the COVID-19 pandemic, amplification-free biosensing concepts including CRISPR-based diagnostics,^13–15^ nanopore readout^16,17^ DNA origami^18,19^ and DNA circuits have been further developed. DNA circuits based on toehold-mediated strand displacement (TMSD) are among the most studied circuits, owing to their versatility, programmability, and amplification-free operation at ambient conditions.^20–24^ While TSMD-based target recycling circuits simplify experimental requirements greatly, they lead to weak signal gain at low target concentrations and have slow kinetics, by TMSD being a statistic process of toehold (TH) formation and branch migration.^21^

Over the past few years, the kinetics of TSMD circuits have been enhanced by using enzymes and proteins.^25–28^ However, for real-world applications and especially for testing in low-income countries, low temperature transport and storage of enzymes and expensive reagents are not an option. To realize nonenzymatic circuits with fast kinetics, elaborate amplification circuits have been employed.^29,30^ Numerous DNA circuits such as catalytic hairpin assembly,^31,32^ hybridization chain reactions^33,34^ and circuit reactions^29,35^ were designed to amplify the signal gained per target. Circuit reactions can utilize TH exchange, which involves partial displacement and spontaneous dissociation of an incumbent strand, opening an allosteric toehold for a so-called fuel sequence, which recycles the target and enhances the reaction kinetics.^29,36^ A study on TH exchange processes has demonstrated the positive effect of the primary TH being one base-pair longer than the allosteric TH on the reaction kinetics.^36^ Other studies have demonstrated the significance of sequence^37,38^ and complementarity of utilized DNA strands, as mismatched base-pairs accelerate or decrease the kinetics of TMSD-based circuits substantially.^39–43^ Implementing a mismatch (MM) in the allosteric TH domain can lead to a synergic effect and enhance the kinetics of TSMD-based circuits significantly. However, it has not yet been explored how the position of MM impacts the kinetics of nonenzymatic diagnostic cascades.

TMSD-based circuits often use colorimetric or fluorescent readout methods and rely on reporter duplexes to monitor their kinetics.^24,36,40^ Alternative readout approaches utilize functionalized gold^44–47^ or MNPs^8,48–55^ as markers. While fluorescence-based assays are affected by background molecules,^56,57^ magnetic measurements are potentially not influenced by cell debris, background macromolecules, and proteins present in complex biological samples.^58,59^ Moreover, magnetic fields are negligibly attenuated in opaque media, thus allowing the detection of nucleic acids directly on unprocessed samples, unlike methods based on visible light. The dynamic magnetic response of MNPs to moderate alternating magnetic fields changes in a highly specific manner upon molecular binding between receptors on MNPs and targets in solution.^46,53,60–63^ The change in particle hydrodynamic size resulting from target recognition, is picked up within a minute with magnetic particle spectroscopy (MPS), a highly sensitive magnetic readout system.^64,65^ Magnetic assays with detection limits of up to 1 fM were developed in recent years, yet only through enzymatic reactions.^66–68^ Recently, we have successfully detected viral genome of SARS-CoV-2 by disassembling clusters of MNPs using a nonenzymatic TMSD-based DNA circuit and reading out the corresponding change in the MPS signal.^35^ With a limit of detection (LoD) of 27 pM after 24 h assay time, the current declustering-based assays are still unable to function at clinical POC settings, where low detection limit and short assay time must be merged.

Here, we propose innovative nonenzymatic declustering-based assays for the detection of nucleic acids in solution by integrating mismatch-assisted toehold exchange (MATE) in magnetic diagnostics cascades. We demonstrate that MATE is a highly efficient means of recycling nucleic acid targets, amplifying the magnetic signal output, and accelerating the assay’s kinetics. Kinetics of target-dependent declustering of magnetic clusters can greatly be tuned by adding fuel strands that recycle the target via TH exchange. Our experimental kinetic studies show the beneficial impact of a mismatched base-pair on the declustering rate and equilibrium state. OxDNA coarse-grained model^69,70^ simulations on the effects of MM position on dissociation/declustering kinetics support the experimental observations. The MM-mediated duplex destabilization enhances the off-rate of the allosteric TH, hybridization of fuel strands, and recycling of the target significantly. Our novel MATE cascades have a LoD of 7.6 pM after a total assay time of 2 h, as determined from MPS measurements. Our work demonstrates that MATE accelerates the kinetics and improves the sensitivity of nonenzymatic magnetic DNA assays drastically, taking a major leap towards translating magnetic nanoparticle-based assays to the clinical POC settings.

## RESULTS AND DISCUSSION

MNP-based diagnostics are nonenzymatic and isothermal,^35,61,71^ and have the potential to be performed directly on unprocessed biological samples, all indispensable features for POC applications. However, the existing MNP-based bioassays are unable to unlock these unique potentials, as their detection limit and time to results do not fulfill rapid testing requirements. So-called declustering-based assays register magnetic signal gain upon dissociation of MNPs from pre-formed magnetic clusters as well as their disintegration into smaller clusters in response to target.^35,61^ Through these two concurrent events, the overall cluster hydrodynamic size decreases and Brownian magnetic relaxation processes become faster on the ensemble level, resulting in the magnetic signal gain at the specifically chosen excitation frequency (see the supporting information (SI) for more details). Over time, we have witnessed that such circuits lead only to partial declustering at low target concentrations (Figure 1a, left panel), providing a low overall signal gain, since the target strands cannot react further after one TMSD event is complete. Here, by integrating TH exchange and mismatch destabilization in magnetic diagnostics cascades, we introduce a novel nonenzymatic declustering-based nucleic acid detection approach named MATE-cascades. TH exchange and base-pair mismatches have individually been employed to improve the kinetics of TMSD-based circuits,^39–41^ yet their combination and usability in magnetic diagnostics circuits are completely unexplored.

**Figure 1.**
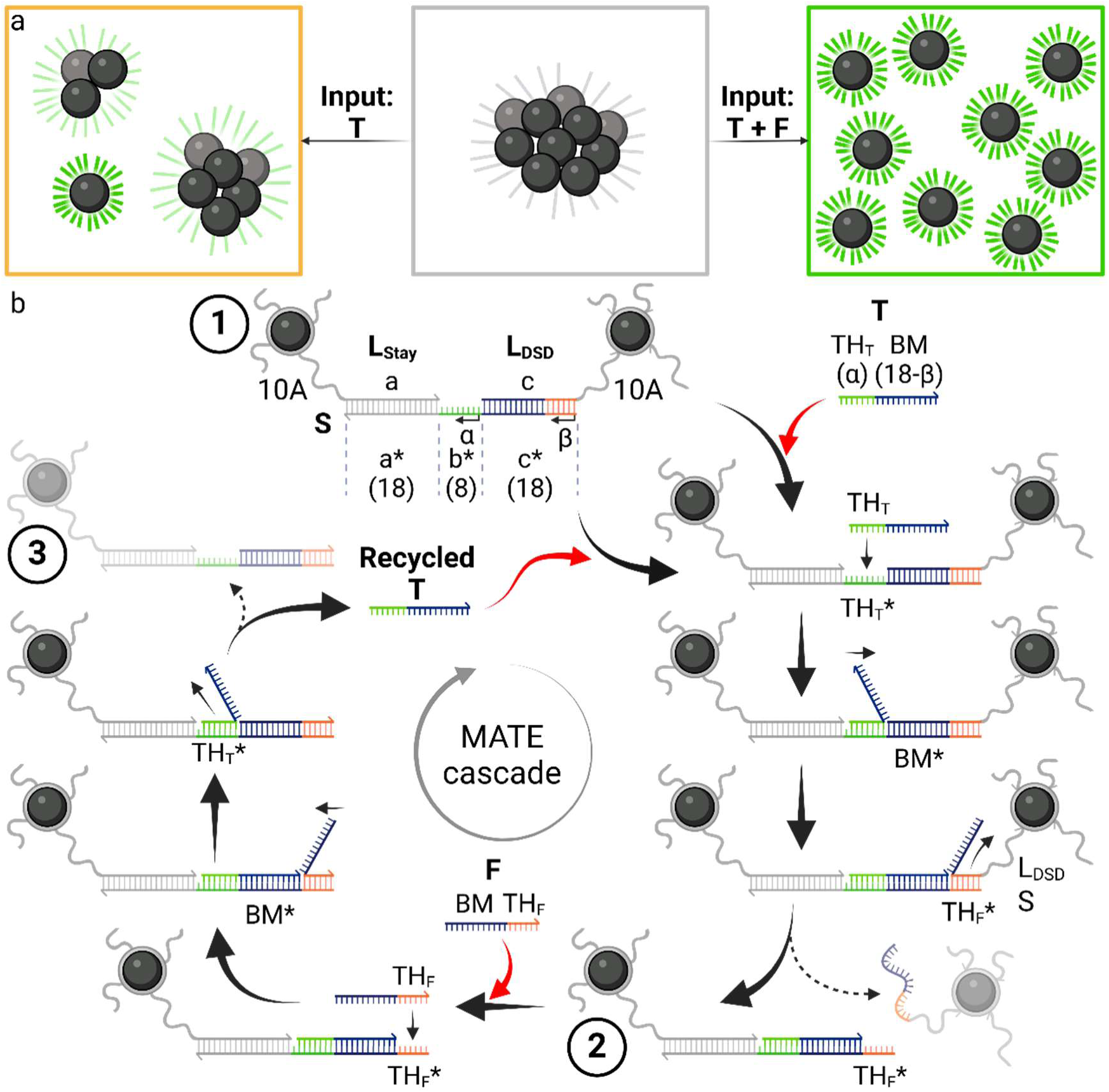
Scheme of MATE diagnostics cascades. (a) Scheme of declustering of magnetic clusters upon addition of target DNA (T) at low concentrations in the absence (middle -> left panel) or in the presence (middle -> right panel) of fuel DNA (F), leading to partial or full declustering, respectively. (b) MATE declustering cascade. First, T displaces L_DSD_. The last β bps of L_DSD_ dissociate spontaneously, resulting in a release of an MNP from magnetic clusters (stage 1 -> 2). Afterwards, TH_F_* becomes accessible for backward TMSD reactions, wherein F docks, branch migrates and releases T (stage 2 -> 3). Next, the recycled T initiates the next cycle, leading to further declustering reactions, thus amplifying the total signal gain at low T concentrations. The magnetic clusters are irregularly shaped 3D structures, here shown as dimers for simplicity.

A typical MATE assay comprises pre-formed magnetic clusters, target (T) and fuel (F) strands. The clusters are formed by combining two types of BNF80-MNPs (see SI for synthesis procedures), one type labelled with label DNA Stay (L_Stay_, domain a) and the other labelled with label DNA DSD (L_DSD_, domain c), with substrate DNA strands (S) (Figure 1b, stage 1, and Tables S1, S2 for the sequences). S consists of domains a* and c*, complementary to the labels, as well as TH domain b* where the toehold TH_T_ of T can attach to. T consists of TH_T_, of α nucleotides, and a branch migration domain (BM). BM is equivalent to domain c, reduced by β bps from the MNP-proximal 3’ end. In a MATE declustering cycle, T displaces L_DSD_ up to the last β base pairs (bps) upon binding to TH_T_* and branch migration. An MNP is then detached from the clusters if the β bps dissociate spontaneously (stage 2). The released MNP and the magnetic ensemble, reduced in hydrodynamic size, increase the magnetic signal output, which can be read out as a gain in magnetic susceptibility values (see Figure S1). Simultaneously, the previously hidden domain TH_F_* is now accessible for F. Strand F attaches to TH_F_* and displaces the domain BM between T and S in a reverse displacement reaction. Upon completion, α base-pairs of TH_T_ dissociate spontaneously from S, completing one MATE cycle by releasing T and feeding it into the next cycle (stage 3).

### Implementing a mismatch in TH_F_ enhances signal gain most if placed at the first bp of spontaneous dissociation

We first studied how the declustering rate depends on the length of β of TH_F_ by varying it from 5 to 8 bp, while keeping α = 7 bp. A systematic reduction in the rate was observed by increasing the length of β compared to full displacement of the L_DSD_ (β = 0 bp, see Figure S2), aligning with the literature.^36,72^ While the declustering with β = 6 bp causes visible magnetic signal change, the overall declustering rate is reduced as a result of slow spontaneous dissociation and thus the signal gain is limited. Consequently, we wondered how implementing a mismatched bp between S and L_DSD_ would increase the declustering rate. We inserted one non-canonical bp at different positions in the spontaneous dissociation domain on S TH_F_*, which locally destabilizes the duplex between L_DSD_ and Sx (MMs are highlighted in red in Figure 2a). The MM identities were chosen based on NUPACK simulations to ensure full hybridization of the MNP-proximal end of the duplex and avoid significant declustering by F in the absence of T (see Table S2 for sequences and estimated energies of the respective duplexes).

**Figure 2.**
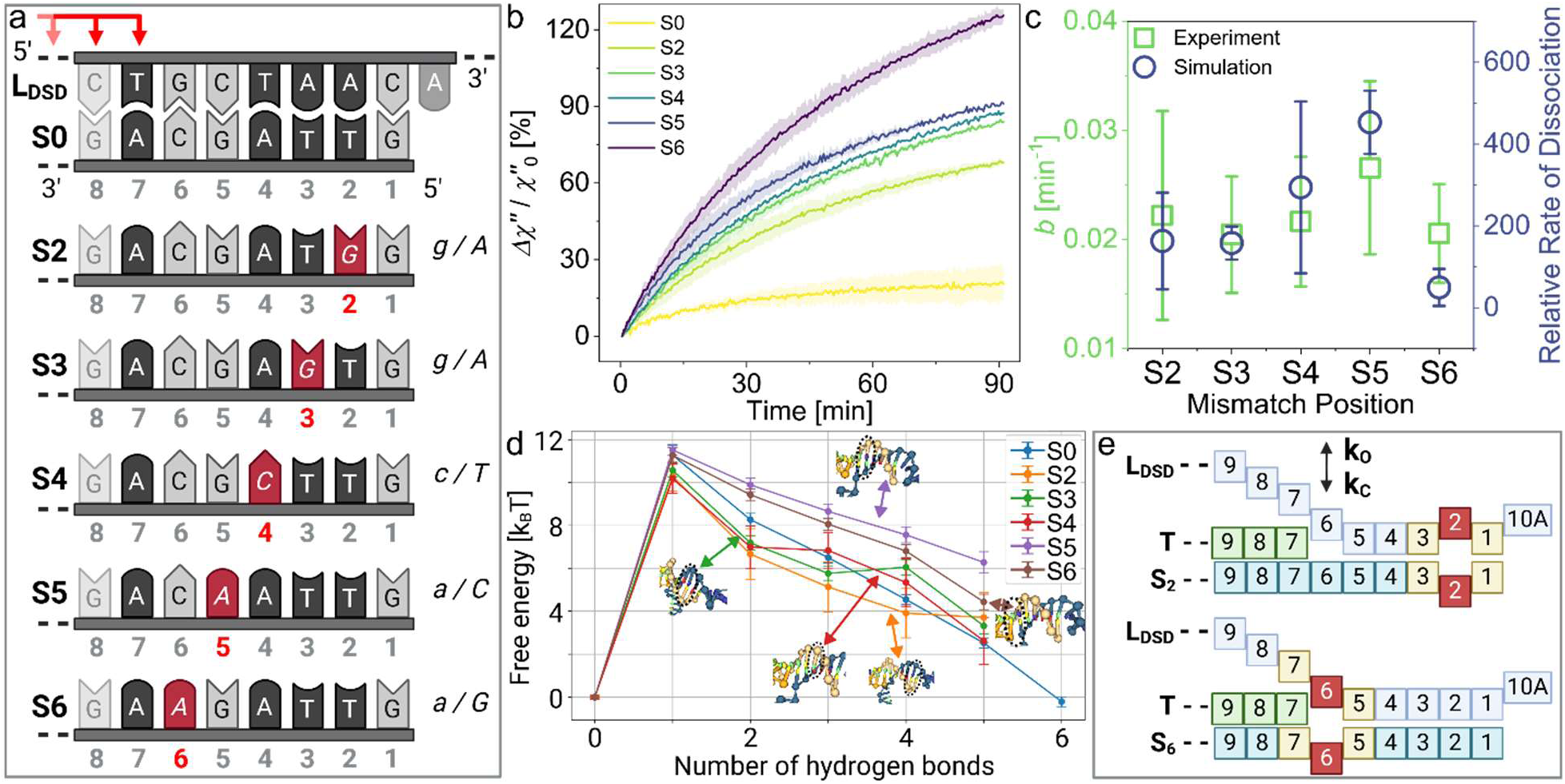
Declustering rate and signal gain as a function of spontaneous dissociation in response to T: (a) Base-pairs at the MNP-proximal end of the L_DSD_:S duplex for full complementarity between S and L_DSD_ (S0) and with MMs at different positions x (Sx), counted from the MNP-proximal end. (b) Relative change in the imaginary part of complex ac magnetic susceptibility *Δχ′′* over time for assays on clusters with different MMs (100 mM Mg^2+^, 4 nM Sx, 1 nM T (α = 7 bp, β = 6 bp), calculated from two independent replica, displayed 1σ error shade. (c) Relative dissociation rate (with respect to no mismatch case S0) as measured in oxDNA forward flux simulation for different mismatch positions, compared to saturation rate *b* fitted to experiments. (d) Visualization of destabilizing effect of MM (red) at position 2 (S2) and 6 (S6), complementary base-pairs are displayed in yellow (destabilized by MM), green (T), and blue (S, L_DSD_). (e) Free-energy profile for strand dissociation for different mismatch positions, as obtained from oxDNA simulations.

To understand how the MM position influences the declustering rate and eventual availability of the toehold TH_F_*, we first performed assays in the absence of F. The kinetics of declustering reactions were monitored by measuring the imaginary part *χ′′* values of complex ac magnetic susceptibility^73,74^ at the excitation frequency of 120 Hz and field strength of 0.5 mT, chosen based on the magnetic measurements at different conditions (see Figure S1). Once declustering occurs, single MNPs are released and the overall hydrodynamic size of the magnetic clusters decreases, both increasing *χ′′*. Looking at the relative change in *χ′′* (eq. S4) measured over 90 min at 1 nM T for two independent replicas, we observed that the gain in magnetic signal strongly depends on the position of MM on S (Figure 2b), implying different declustering rates (Figure 2c). In case of a MM-free duplex (S0), the reaction saturates quickly with only a small signal gain, indicating inhibited declustering at S0. The closer the MM is to the last base-pair between T and S (bp 7), the more declustering reactions occur.

To quantify the declustering processes, we fitted the experimental data to an exponential function given by

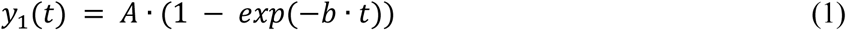

with time *t* in min, saturation rate *b* in min^−1^ and total signal gain *A* in % (see Table S3 for all values). Parameter *A* increases by placing the MM closer to where T ends, with the cases S6 and S0 showing the highest and lowest values, respectively. The averaged saturation rate *b* shows an intriguing and rather unexpected trend (Figure 2c). It increases from S2 to S5 and then drops for S6. S5 shows the highest *b* value of 0.027 min^−1^. While the circuits with MM S6 reach their thermodynamic equilibrium after a longer time with *b* = 0.021 min^−1^, its *A* = 147.9 % value is 7.1-fold higher than the case of full complementarity S0 (*A* = 20.9, see Table S3 for fit parameters prior to averaging).

To further investigate the mechanism of spontaneous dissociation of L_DSD_ (the step immediately preceding stage 2 shown in Figure 1b), we used oxDNA simulations to gain insight into the process. We obtained free-energy profiles as a function of number of base-pairs formed for different positions of the MM between the substrate and the dissociating strands. The profiles (Figure 2d) show that the MM position 5 (S5) is the most destabilizing, as seen by the relative free energy difference between the fully unbound (0 bp) and fully bound strand (5 bp). Other MM positions were estimated from the profile to be about 1 to 4 kT more stable in the bound state than the S5 one. This was also confirmed with our forward flux sampling (see Figure 2c), where the dissociation with MM in position 5 was the fastest, aligning well with our experimentally derived saturation rates *b*. Thus, our dissociation study illustrates how mismatch position can be also used to fine-tune spontaneous dissociation kinetics in strand displacement cascades.

Relevant for our sensing application is *A* after certain run-time. If the system reaches its thermodynamic equilibrium quickly, but does not yield a significant change in signal, it limits the potential LoD and is therefore not suitable for the detection of low numbers of target DNA. Our results clearly show that the total signal gain (*A*) and the correlating overall declustering with spontaneous dissociation of 6 bps are increased if a MM is positioned adjacent to the last bp of the hybridized T. Two mechanisms may explain why positioning a MM further away from the BM domain reduces the overall declustering. One explanation might be that the TH_F_ domain detaches from Sx and reattaches to a free TH_F_* domain on Sx. As a result, MNPs are always seen in the clustered state and thus no magnetic signal *Δχ′′* is gained. To test this hypothesis, we added T with blocker strands to the cascade, which would attach to the TH_F_* and TH_F_ domains once those are accessible after spontaneous dissociation of L_DSD_. We observed no significant change in the declustering behavior (see Figure S4 for data and Table S4 for sequences). Therefore, we argue that a more plausible reason for limited declustering behavior is the difference in initial detachment of TH_F_ once T has finished the BM step. The TH exchange reaction is a stochastic process based on forward and backward steps of each base-pair involved. Once T is fully hybridized to S, the detachment of TH_F_ depends on the opening *k*_O_ and closing *k*_C_ rates of the last 6 bps of L_DSD_. If bp 6 opens, the remaining 5 bps are less stable and are increasingly prone to spontaneous dissociation. Conversely, if bp 6 is connected to S, L_DSD_ can initiate a reverse TMSD reaction and displace T. This will, in turn, limit the declustering processes and the magnetic signal gain *Δχ′′*, as schematically shown in Figure 2e with the MM colored in red and the adjacent, destabilized nearest neighbors (NNs) in yellow. Both *k*_O_ and *k*_C_ depend on the local energy landscape of the duplex. The presence of a MM destabilizes its immediate location and the two NNs and deforms the helical structure for up to few neighboring bps.^75^ In the proximity of a MM, the opening rate *k*_O_ outweighs the closing rate *k*_C_, since the formation and enclosing of a MM is energetically unfavorable.^39,43,76,77^ While a MM at position 2 (S2) enhances *k*_O_ of TH_F_ once the detaching branch has reached the last bps (yellow and red), it has marginal influence on *k*_O_ at position 6 (blue). Therefore, the MM in S2 inhibits the backwards pathway only minimally, due to the complementary domain between TH and L_DSD_ being reduced to 4 consecutive bps. The closer the MM is shifted to position 6, the earlier the undesired backwards path by L_DSD_ is hindered. At position 6 (S6), an energetic barrier has to be overcome to form a bp at the position of the MM (red)^76,78^ and be closed long enough for the bp between T and S at position 7 (green) to initiate a reverse TMSD reaction (yellow). The probability for this event to take place is significantly lower than for TH_F_ to detach once T has completed its branch migration. Additionally, the forward branch migration of the last bps of T is increased due to the local destabilization caused by the MM at position 6. One further influential factor could be the distance of the MM to the duplex end. A MM may have difficulties to form a closed base-pair if it is enclosed by numerous bps that are forcing a helix-structure, while a lower number of bps adjacent to a MM, being closer to the duplex end, can deviate easier from the helix-structure to adapt to the MM deformation.

### Recycling efficiency in magnetic cascades increases with the length of allosteric TH_F_

Moving forwards with our investigations, we realized that adding 100 mM Mg^2+^ to the cascade improves the declustering up to 17-fold by enhancing hybridization of TH_T_, similar to what has been reported by Yang et al.^79^ Importantly, Mg^2+^does not impede the spontaneous dissociation of TH_F_ (Figure S5). In TH exchange-based cascades, the difference in bp of TH_T_ and TH_F_ (α − β) plays as equally important role as the length of each TH domain,^36^ yet it is completely unknown how this reflects in magnetic signal amplification cascades like ours. To shed light on this aspect, we varied the length α of TH_T_ from 5 to 7 bp, while keeping β = 6 bp and measured *Δχ′′* in the presence of T. The overall signal change increases with (α − β) for both 0 and 100 mM Mg^2+^ (Figure S6). Having more bps in TH_T_ (α – β = +1 bp) enhances the TH formation and favors the hybridization of T over L_DSD_ to S thermodynamically, both effects improving the subsequent declustering rate. To this end, we opted to continue with a TH_T_ length of α = 7 bp to enable fast and reliable opening of TH_F_*.

We then sought to understand how efficiently our MATE-cascade can recycle T and feed it back into the next declustering cycle if the input F was added. F binds to TH_F_* only if it is available upon the completion of declustering (stage 2 in Figure 1b). Hence, adding F to the MATE-cascade recycles T in a reverse TMSD pathway (stage 2 → stage 3), feeds it back into the next cycle, and completes one signal amplification cycle. We tested two different lengths β of TH_F_ in the presence and absence of a MM in TH_F_ (Figure S7). Over three independent sets of experiments, β = 6 bp showed significantly higher recycling capabilities compared to β = 5 bp. In this situation, the reverse reaction is offered a longer, more stable TH, enabling F to initiate the recycling of T more efficiently. Another advantage lies in the nature of TH exchange, where upon detachment of the α = 7 bp of TH_T_, 2 bps are lost for β = 5 bp (β – α = −2 bp), while the duplex is deprived of only 1 bp for β = 6 bp (β – α = −1 bp). These two factors align to enhance the reverse TMSD reaction and the related recycling of the target DNA. These effects are even stronger in the presence of a MM in TH_F_, showcasing the importance of a longer TH_F_ domain for enhanced recycling.

### MATE increases target recycling by 14-fold

These findings motivated us to look into how adding F changes the signal gain for all the mismatch positions for α = 7 bp and β = 6 bp (Figure 3a). The trend of *Δχ′′* over 90 min between the different mismatch locations increases non-chronologically via 0 ≪ 4 ≤ 2 < 5 ≪ 3 < 6. The magnetic signal for an assay with complete MATE-amplification is enhanced by 5.2-fold from Δχ′′ = 34 % for S0 to Δχ′′ = 176 % for S2. Although F is present in excess (100 nM F on 4 nM S and 1 nM T), the disassembly of S0-based, MM-free clusters is significantly restrained due to the slow and limited opening of the allosteric TH_F_*. The *Δχ′′* increases most for S6 by 14.4-fold to 490 %. The cascade of S0-based clusters reaches saturation after 1.5-2 h of monitoring, while *Δχ′′* of the samples with mismatched substrates still increases significantly after 1.5 h, indicating that the declustering and signal amplification are still significantly in progress. This behavior is based on the cooperation of the spontaneous dissociation of TH_F_ after TMSD of T and leakage, which is declustering caused by F invading at breathing bps at the MM location.

**Figure 3.**
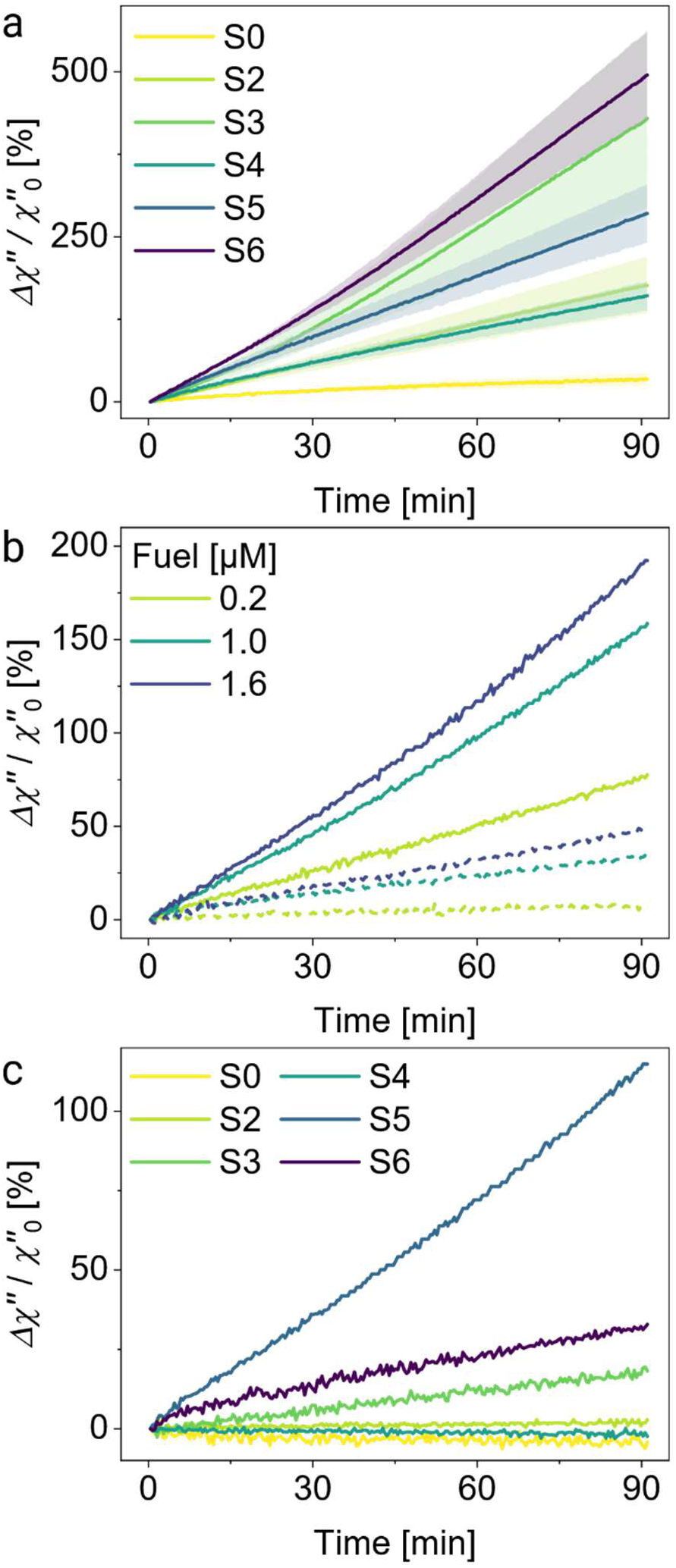
Magnetic signal amplification via recycling of T by F. (a) *Δχ′′* for all MM positions Sx (1 nM T (α = 7 bp, β = 6 bp), 100 nM Fx, complementary to Sx (4 nM)). The curves are the average of three independent sets of samples (1σ error shades). (b) *Δχ′′* on S6 for increasing Fuel concentrations with 0.25 nM (solid) and without T (dashed). (c) *Δχ′′* for different MM positions of Sx at 1 µM Fx in the absence of T (leakage). All measurements contained 100 mM Mg^2+^.

We next explored to what extent the recycling of the target and its reuse in subsequent declustering reactions can be accelerated by concentration-driven TMSD.^21^ To do so, we varied the F concentration from 4 nM (F:S 1:1) to 100 nM (F:S 25:1) and looked at how the magnetic signal changes accordingly (Figure S8). Due to the loss of a bp in the reverse TMSD path (β - α = 6 - 7 = −1 bp), the hybridization product SF is slightly less favorable than ST, requiring oversaturation of F over S. We measured *Δχ′′* for clusters with a MM at position 6 (S6) at three F concentrations (Figure 3b) in the presence (solid lines) and absence (dashed lines) of 0.25 nM target. After 90 min, *Δχ′′* increases from 77 to 192 % and 9 to 49 % by increasing F concentration from 0.2 to 1.6 µM in the presence and absence of T, respectively. Admittedly, we observed some declustering in the absence of T, an effect known as leakage (dashed lines, Figure 3b), which increases with the F concentration. The leakage is presumably due to enhanced breathing of the domain TH_F_ resulting from duplex destabilization by the MM.

### Highest opening rate of TH_F_ by T does not entail highest leakage

Moving towards establishing an assay workflow using MATE, it is important to understand the origin of the leakage. To shed light on this critical topic, we first looked at how the leakage depends on the MM position at high F concentrations, as different MMs destabilize the duplex between L_DSD_ and Sx differently. We recorded *Δχ′′* for all MMs by adding 1 μM Fx to different MATE cascades (Figure 3c) in the absence of T. We observed −5.3, −2.3, and +1.8 % change in *Δχ′′* for S0, S4, and S2, respectively, over 90 min incubation time, indicating no significant leakage for these designs. The situation for other designs is quite different. The S3 and S6 show 19.1 and 31.2 % signal change leakage. Most significant is the leakage for S5 with 114.8 % increase in *Δχ′′*. We observed a similar trend in leakage for the MMs studied for other concentrations of F, with the effect increasing with concentration (Figure S9).

To understand these results better, we calculated the free energy Δ*G* values (Table 1) of the DNA complexes in clusters (stage 1 in Figure 1b) via NUPACK. We found that the difference in Δ*G* increases by placing the MM further away from the MNP proximal end and climaxes for S5, matching the experimental observations (Figure 3c). Based on the differences in Δ*G*, we expected S3 to show similar leakage behavior as S2 and S4, which is not the case. Inspecting the NN of the mismatched bp, we notice that the MM at S3 is surrounded by two A-T pairs, which are more likely to breathe than G-C pairs.^75^ In S2 and S4, the MM is enclosed between one A-T and one G-C pair. While the MMs at position 5 and 6 are adjacent to one G-C NN, the mismatched base-pair replaces a G-C pair, instead of an A-T pair (as for S2 and S4), and reduces the GC-content in the THF domain. Therefore, we hypothesize that not only the base-pairs directly adjacent to the MM are relevant for the duplex stability, but bases further away from the MM also play an important role. The leakage for S5 may additionally be higher due to the MM identity chosen in this study. Oliveira et al. observed A-C pairs lacking strong hydrogen bonds^42^ and Rossetti et al. determined A-C to show the highest breathing portions of the MMs studied in our work.^75^ Interestingly, the most probable structure which NUPACK calculated for S2 shows only a probability of 60 % for the last bp to be closed (see Figure S10). This would entail that the last one to two base-pairs would be present in open constellation for nearly half of the time, which would lead to significant leakage. Oliveira et al. reported that the MM A-G within the sequence AaC/TgG exhibits a double hydrogen bond and is therefore exceptionally stable,^42^ which may explain the observed low leakage. We acknowledge the influence of the identity of the MM as well as the NN on the stability of the duplex.^42,80^ The sheer number of MM-NN combinations go beyond the scope and capability of this research.

**Table 1.** Sequences of the MM and its NN: mismatched base at position x as lower-case letter, Δ*G* simulated with NUPACK for 150 mM Na^+^, 100 mM Mg^2+^, 5 nM Oligos (L_DSD_, L_Stay_, Sx) at 25 °C.

| Substrate<br>Sx | MM + NN<br>identities<br>x±1 | $\Delta G$<br>[kcal/mol] | $\Delta G(S0) - \Delta G(Sx)$<br>[kcal/mol] |
| --- | --- | --- | --- |
| S0 | - | -59.77 | ±0 |
| S2 | AAC/TgG | -57.14 | +2.63 |
| S3 | TAA/AgT | -56.39 | +3.38 |
| S4 | CTA/GcT | -56.08 | +3.69 |
| S5 | GCT/CaA | -54.32 | +5.45 |
| S6 | TGC/AaG | -54.93 | +4.84 |

### Local duplex stabilization around MM regulates leakage and spontaneous dissociation of TH_F_

The peculiar correlation between the MM position and leakage motivated us to explore how the leakage is modulated by duplex stabilization around MM, with a focus on S6, as it shows the highest spontaneous dissociation of the TH_F_ domain and comparatively low leakage. To do so, we modified S6 in the TH_F_ domain in two specific ways. First, we replaced a single nucleotide with a locked nucleic acid (LNA) at different neighboring positions without changing the base identity (Table 2) and looked at how the stabilizing nature of the LNA^81,82^ influences the leakage and target-catalyzed declustering rates. Further, we also modified the last two base-pairs from the MNP proximal end by inserting a CG-clamp (see Table 2 and S5). We compared the leakage by monitoring *Δχ′′* over 90 min by adding only F to the MATE cascade (Figure 4a). Remarkably, the cascade shows higher change in *Δχ′′* and therefore faster declustering for the CG-variation than for the basic version, suggesting that the leakage does not stem from the duplex end and that the increased CG-content in THF (33.3 → 50 %) facilitates hybridization of F to the TH_F_* domain. Placing an LNA at position 2 and 5 stabilizes the L_DSD_:S duplex structure only slightly and reduces the leakage marginally. The behavior is very different when the LNA is at position 7, where a significant drop in the leakage was observed (Table 2). Our results strongly suggest that the leakage stems mainly from the A-T pair at position 7. We, therefore, hypothesize that the MM destabilizes its NN asymmetrically, with the destabilization and breathing of the A-G MM at position 6 being more prominent towards the A-T bp than towards the G-C NN. Therefore, employing an LNA at position 7 inhibits the leakage by F most effectively.

**Table 2.** Magnetic signal gain *Δχ′′* over 90 minutes for different TH_F_ modifications. Varying substrate stability from the basic concept (S6) by addition of a LNA yz (y: position from MNP-proximal duplex end, z: base identity) marked with {…}, as well as a CG-clamp. MM as lowercase letter. *Δχ′′* after 90 minutes of declustering with F (leakage), with T (no recycling) and with fully functional MATE recycling and amplification. Ratio of *Δχ′′* in leakage relative to recycling.

| Sequence variation | Last 8 nt of S6 (3'→5') | Leakage [%]<br>1 μM F | No recycling [%]<br>1 nM T | Recycling [%]<br>0.25 nM T, 1 μM F | Leakage / recycling [%] |
| --- | --- | --- | --- | --- | --- |
| Basic | ... GAaGATTG | 35.5 | 155.4 | 207.4 | 17.11 |
| CG | ... GAaGATGC | 54.2 | 150.0 | 228.1 | 23.76 |
| LNA 2T | ... GAaGAT{T}G | 30.5 | 159.1 | 157.1 | 19.41 |
| LNA 5G | ... GAa{G}ATTG | 29.2 | 114.8 | 126.9 | 23.01 |
| LNA 7A | ... G{A}aGATTG | 15.3 | 120.9 | 121.6 | 12.58 |

**Figure 4.**
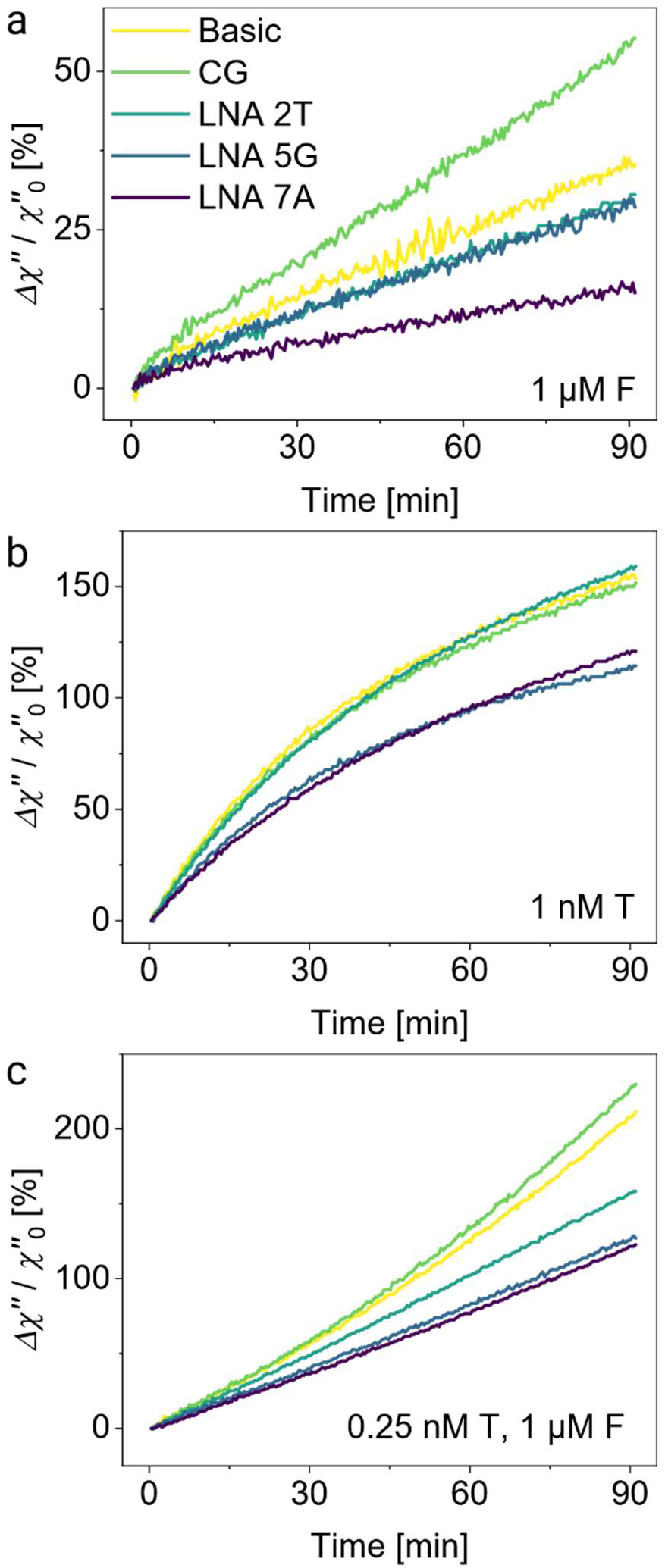
Reducing leakage with locked nucleic acids. Changes in magnetic signal over 90 min for (a) Leakage-based declustering (1 µM F) in the absence of T, (b) declustering at 1 nM T in the absence of F, and (c) the whole MATE cascade at 0.25 nM T and 1 µM F.

Next, we investigated the influence of the duplex modifications on the spontaneous dissociation, since this effect is at the core of MATE cascades. We observed no significant change in Δχ′′ for the CG-variation and the LNA at position 2 (Figure 4b). The declustering decreases for the LNAs in position 5 and 7. Therefore, we argue that the last two bps do not impact the spontaneous dissociation of TH_F_ in contrast to the NNs of the MM. The LNAs at position 5 and 7 reduce the opening rate of TH_F_* noticeably as the incumbent is stabilized around the MM.

Highly relevant for our magnetic DNA cascade is the full MATE cascade amplification, where T is recycled back into the circuit by F (Figure 4c). A minimal increase in *Δχ′′* can be observed for the CG-variant (Table 3; *δ*(Δχ′′/χ′′_o_) = +9.9 %, calculated using eq. S6), which is plausibly due to increased leakage (+52.7 %) rather than a better functioning cascade. The magnetic signal gain *Δχ′′* and its underlying declustering efficiency are reduced for all tested LNAs due to the reduction in leakage and spontaneous dissociation. The LNA in position 7 is the only variation that features a reduction in leakage to recycling ratio (Table 2) and is therefore the most favorable S6 variant for diagnostic applications using the MATE cascade.

### MATE-based cascades depend linearly on target concentration

By combining all the optimal conditions established so far, here, we work towards realizing MATE cascades for diagnostics. Up to this stage, we primarily looked at how a MATE cascade responds to single base-pair variations. Here, we asked how a full MATE cascade functions if F and T concentrations are varied over a broad range. First, we varied the F concentration from 0 to 4 μM at 0.25 nM T and monitored changes in *χ′′* (Figure 5a). The comparably low target concentration was chosen to simulate the performance of our assay and show the relevance of MATE-based amplification for clinical samples with a low target DNA concentration. For all samples, we observe a linear increase in *χ′′*, except for the case at 0 μM F, which saturates within the recorded time due to no T recycling and signal amplification. By increasing the fuel concentration, a much greater change in signal *χ′′* after a shorter time was observed, with the steepest rise for 4 µM F. To estimate the reaction rate, we fitted the measured data to a simple linear function given by

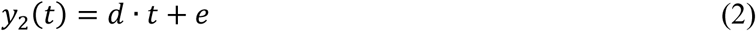

**Figure 5.**
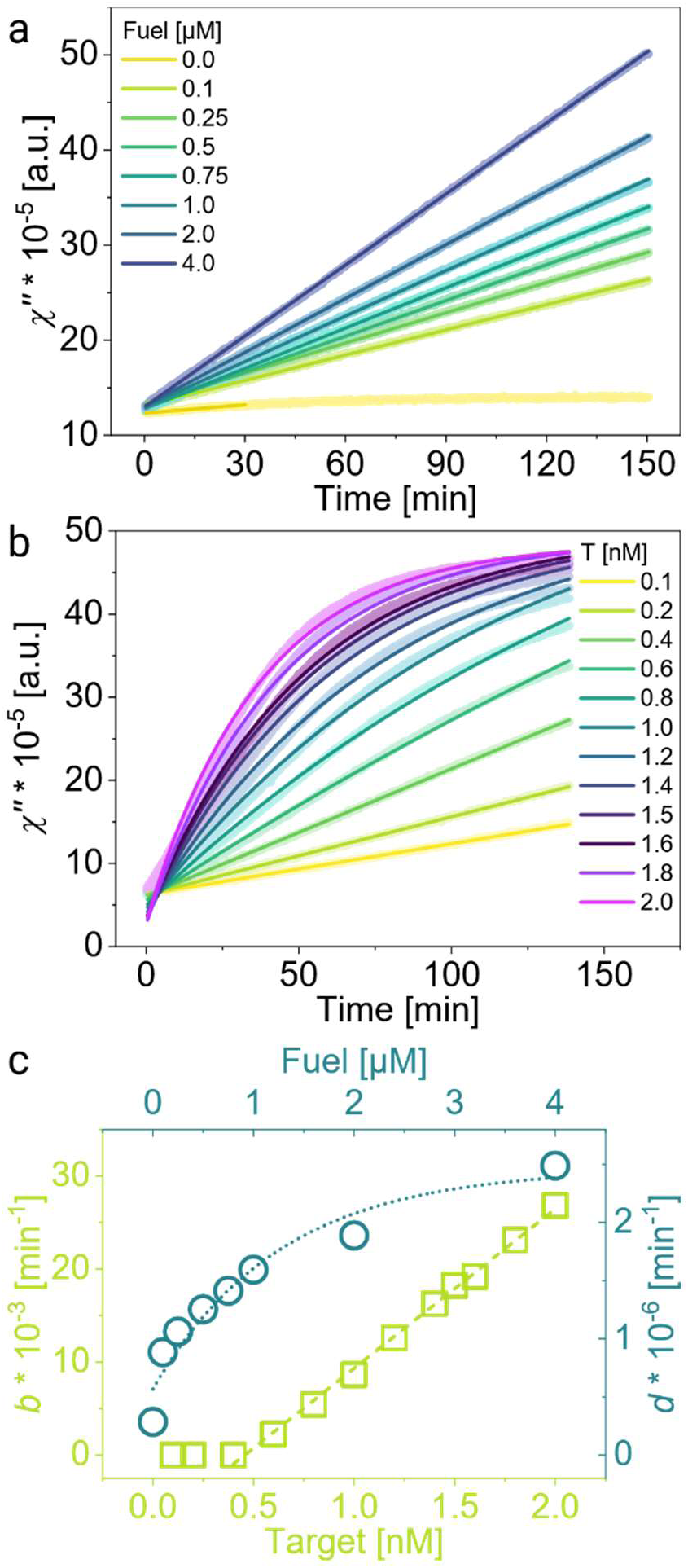
Concentration dependent MATE signal amplification Input DNA concentration dependent measurements. (a) *χ′′* measured at different F concentrations at 0.25 nM T, solid lines were fit with eq. 2. (b) *χ′′* at different T concentrations at 4 µM F6, solid lines were fit with eq. 3. (c) Parameters *b* and *d* from fit functions over respective concentration, dashed lines were plotted via eq. 2 (green, target series) and 3 (blue, fuel series). All experiments were executed with α = 7 bp, β = 6 bp, S6, LNA variation in position 7, 100 mM Mg^2+^. All clusters within one panel are from one preparation batch, enabling plotting magnetic susceptibility *χ′′* as recorded.

with *d* the slope in min^−1^ and *e* the offset in a.u., compensating for minor particle concentration differences. The curve at 0 μM F was fitted only up to the first 30 min where *χ′′* increased linearly.

In another set of experiments, we recorded changes in *χ′′* by varying the T concentration from 0.1 to 2 nM at a constant F of 4 μM (Figure 5b). The kinetics of declustering and signal amplification seem very different compared to when the F concentration was increased. The *χ′′* increases faster with [T]. Higher T concentrations lead to higher opening rates of TH_F_* and efficient hybridization of F strands to fully accessible allosteric THs. This in turn leads to more efficient target recycling and a complete declustering of magnetic clusters (Figure 1a). At [T] > 0.6 nM, *χ′′* rises exponentially and saturates within the measured time, yet the saturation onset shifts to earlier times by increasing the T concentration. Interestingly, we can observe a slight phase lag for the investigated concentrations (Figure S11), which could indicate a cooperativity between F and T on disassembling our magnetic clusters. We fitted these curves to a modified version of equation 1:

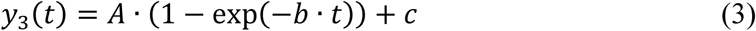

with the parameter *c* the offset of the non-normalized data *χ′′* (in a.u.).

The declustering rates, *d* and *b*, obtained from these two concentration-dependent series reveal interesting features (Figure 5c). By increasing [F] up to 16 000-fold of [T], *d* increases exponentially as a function of [F]. At first, upon adjusting [F] from 0 to 0.1 μM, *d* soars due to the recycling of the low number of T strands. Next, it saturates at high [F], strongly suggesting that there is an upper limit for the impact of F on the declustering kinetics, since the MATE cascade is catalyzed by T. In the experimental series, in which [T] was varied at 4 μM F, very different kinetics were observed (Figure 5b). A saturation in signal can be seen for [T] > ∼ 1 nM, which indicates that no more single MNPs can be generated in the cascade. The rates *b* obtained for this data set increase linearly with [T] at 4 μM F, when the first two data points were excluded, since the MATE cascade is catalyzed by T.

To better understand these data, we broke down the MATE cascade into two reactions at elementary steps:

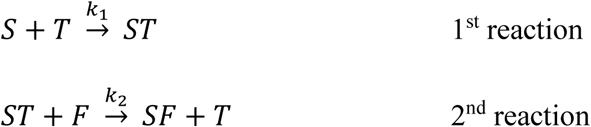

with ST the declustering product (i.e. MNPs), SF the product after recycling of T, and *k*_1_ and *k*_2_ second-order rate constants. The 1^st^ reaction corresponds to declustering and magnetic signal gain, where the cascade transits from stage 1 to 2 (Figure 1b). The 2^nd^ reaction stands for recycling of T by F as the cascade moves from stage 2 to 3. Considering these two reactions, we can tell that the magnetic signal gain comes from the formation of ST in the 1^st^ reaction. In other words, the consumption of ST in the 2^nd^ reaction does not directly change the signal, since no MNP is released. Though consumption of ST does not increase the magnetic signal, the 2^nd^ reaction recycles T strands which will be consumed in 1^st^ reaction pathways, influencing *k*_1_. As we do not have any means to separately determine *k*_1_ and *k*_2_, we hence discuss the reaction rates qualitatively. In the experiments in which we varied [F] at 0.25 nM T, we observed no signal saturation (Figure 5a), indicating a continuous generation of ST over the monitored time. In combination with the linear dependence of kinetic rate *b* on [T] this means that the 1^st^ reaction is the bottle neck of the MATE cascades, wherein at low [T] limited amount of ST is produced and a low number of TH_F_* is catalyzed.

### MATE diagnostic cascades have 3.6-fold higher sensitivity in 12-fold shorter assay time than existing circuits

We then wondered how sensitive and rapid a diagnostics assay based on MATE would be. For these assays, we used MPS, a highly sensitive technique. By doing assays at different T concentrations, we determined the LoD of our MATE-based diagnostic cascade and compared it to our previous magnetic DNA assay.^35^ We combined clusters with 4 nM S6 and the LNA at position 7 with 4 μM F, 100 mM Mg^2+^ and increasing target concentrations (0 nM to 2 nM). The samples were incubated for 2 h at 25 °C and then measured with our custom, bench-top ImmunoMPS system.^65^ The *HR*_53_ of magnetic clusters (Figure 6) starts at a low value (0.1265) at 0 nM T and increases with T as a result of MNPs dissociating from the clusters. Once the clusters are fully disassembled into single MNPs, the *HR*_53_ saturates at *HR*_53_ = 0.1909. Since there was some leakage in the presence of F in prior experiments, the control sample without T contains the same amount of F as the other samples and defines the cut-off value via the 3σ-criterion (light grey dashed line). The *HR*_53_ data was fitted with equation 3. The intersection of the fit curve (dotted line) and the cut-off line determines the theoretical LoD of our assay to be 7.6 pM (≈ 0.57 fmol) after just 2 h.

**Figure 6.**
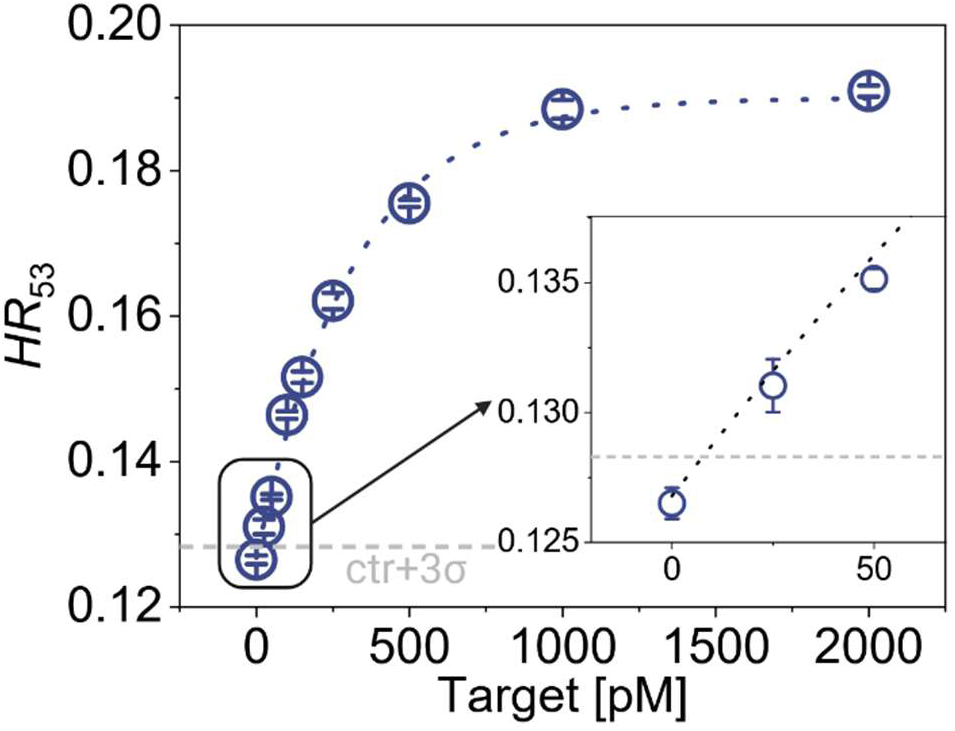
LoD determination for MATE magnetic cascades. Target concentration series measured with custom-made magnetic particle spectroscopy (MPS) after 2 h incubation at 4 µM F, α = 7 bp, β = 6 bp, S6, LNA variation in position 7, 100 mM Mg^2+^. The results are the average of two independent sets of samples.

## CONCLUSIONS

In this work, we proposed mismatch-assisted TH exchange (MATE), a novel concept for next-generation magnetic diagnostics cascade. The MATE cascade offers sensitive, simple, nonenzymatic, isothermal, amplification-free, and rapid diagnostics of nucleic acid sequences in solution. Here, we studied the kinetics of the cascade thoroughly by dissecting it into allosteric TH_F_ generation and target recycling. We showed that a longer allosteric TH_F_ leads to more efficient target recycling and higher signal gain. However, this initially reduces the rate and probability of the spontaneous dissociation of the allosteric TH_F_. By implementing a mismatched bp in TH_F_, the reaction kinetics were enhanced significantly. The spontaneous dissociation rate was enhanced by factor of ∼ 7 for a MM adjacent to the last bp of the invading target, leaving 5 consecutive bps to detach spontaneously.

One major limitation of previous declustering-based magnetic assays is slow TH hybridization. We solved this issue by adding divalent Mg^2+^ ions to the cascade, which accelerated the assay by 17-fold, via stabilizing TH_T_ but not inhibiting the detachment of TH_F_. We improved the assay speed further via concentration driven TMSD by adding more fuel strands and thereby gaining more signal at low target concentrations. Simultaneously, we observed MM-dependent leakage. Single LNA variations were inserted to restrict the leakage-based declustering, resulting in a favorable balance between the unwanted leakage and desired acceleration of the cascade. Our results show that the leakage in MATE cascades depends on the local design of the duplex in terms of the MM identity as well as its surrounding sequence. By performing MATE assays with our bench-top MPS and moving towards establishing simple and rapid assays, we determined the MATE cascade LoD to be 7.6 pM after a total assay time of 2 h. Putting these results into the perspective and comparing them with our previous circuit design,^35^ we witnessed that the novel MATE cascade improved the assay sensitivity by 3.6-fold and the assay time by 12-fold. Looking into the future, we envision that the sensitivity and assay time of MATE can further be improved by incorporating a two-factor amplification design, which combines MATE-based target recycling with the release of an amplification strand, similar to our previous work.^35^

Our study demonstrates that declustering-based magnetic assays benefit substantially from engineering DNA circuits and integrating mismatch and toehold exchange concepts into the magnetic signal amplification cascade. The MATE diagnostics cascade advances magnetic-based assays towards their real-world POC application by offering highly sensitive and rapid assays in a nonenzymatic and isothermal fashion.

## ASSOCIATED CONTENT

Experimental details as mentioned in the text including: chemicals, DNA sequences, sample preparation protocols, assay protocols, oxDNA simulations, and characterization techniques. The SI includes Figure S1 until Figure S11, as well as Tables S1 to S5.

## AUTHOR INFORMATION

### Author Contributions

A.L. and R.S. conceived the research idea. R.S. designed the research, designed the cascades, prepared and characterized clusters, performed the assays, analyzed the data, prepared the figures, and wrote the first draft of the manuscript. J.E. performed oxDNA simulations and analyzed the data. F.W. and T.V. designed and built the MPS spectrometer, M.S. provided resources. P.S. performed and analyzed oxDNA simulations. A.L. designed the research, supervised the study, analyzed the data, and wrote the manuscript. All authors have given approval to the final version of the manuscript.

## NOTES

The authors claim no conflict of interest.

## ACKNOWLEDGMENT

This work is supported by German Science Foundation (DFG) research grants (LA 4923/3-1, VI 892/4-1), Junior Research Group “Metrology4life”, and the Add-on fellowship of Joachim Herz Foundation (R.S.). P.S. acknowledges financial support from National Science Foundation under Grant No 2211794. We thank Miss Petra Schmidt (TU Braunschweig) for ICP-OES measurements. We also thank Dr. Sharif Najafi Shirtari (Kiel University) for fruitful discussions on the reaction kinetics.

## ABBREVIATIONS

## TOC

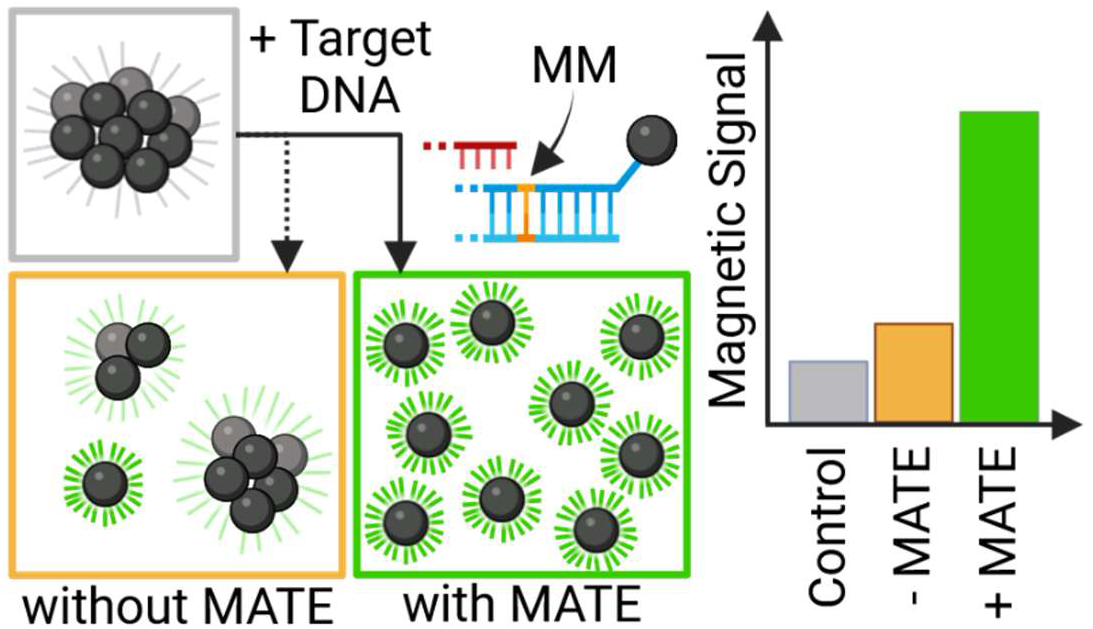

## REFERENCES

(1) Mach, K. E.; Kaushik, A. M.; Hsieh, K.; Wong, P. K.; Wang, T. H.; Liao, J. C. Optimizing Peptide Nucleic Acid Probes for Hybridization-Based Detection and Identification of Bacterial Pathogens. Analyst 2019, 144 (5), 1565–1574. 10.1039/c8an02194e.

(2) Huete-Stauffer, T. M.; Arandia-Gorostidi, N.; Alonso-Sáez, L.; Morán, X. A. G. Experimental Warming Decreases the Average Size and Nucleic Acid Content of Marine Bacterial Communities. Front. Microbiol. 2016, 7 (MAY). 10.3389/fmicb.2016.00730.

(3) Zhou, L.; Chandrasekaran, A. R.; Punnoose, J. A.; Bonenfant, G.; Charles, S.; Levchenko, O.; Badu, P.; Cavaliere, C.; Pager, C. T.; Halvorsen, K. Programmable Low-Cost DNA-Based Platform for Viral RNA Detection; 2020; Vol. 6. https://www.science.org.

(4) Gong, S.; Zhang, S.; Wang, X.; Li, J.; Pan, W.; Li, N.; Tang, B. Strand Displacement Amplification Assisted CRISPR-Cas12a Strategy for Colorimetric Analysis of Viral Nucleic Acid. Anal. Chem. 2021, 93 (45), 15216–15223. 10.1021/acs.analchem.1c04133.

(5) Xu, Z.; Chen, D.; Li, T.; Yan, J.; Zhu, J.; He, T.; Hu, R.; Li, Y.; Yang, Y.; Liu, M. Microfluidic Space Coding for Multiplexed Nucleic Acid Detection via CRISPR-Cas12a and Recombinase Polymerase Amplification. Nat. Commun. 2022, 13 (1). 10.1038/s41467-022-34086-y.

(6) Choi, H. K.; Yoon, J. Nanotechnology-Assisted Biosensors for the Detection of Viral Nucleic Acids: An Overview. Biosensors. MDPI February 1, 2023. 10.3390/bios13020208.

(7) Metcalf, G. A. D. MicroRNAs: Circulating Biomarkers for the Early Detection of Imperceptible Cancers via Biosensor and Machine-Learning Advances. Oncogene 2024, 43 (28), 2135–2142. 10.1038/s41388-024-03076-3.

(8) Tavallaie, R.; McCarroll, J.; Le Grand, M.; Ariotti, N.; Schuhmann, W.; Bakker, E.; Tilley, R. D.; Hibbert, D. B.; Kavallaris, M.; Gooding, J. J. Nucleic Acid Hybridization on an Electrically Reconfigurable Network of Gold-Coated Magnetic Nanoparticles Enables MicroRNA Detection in Blood. Nat. Nanotechnol. 2018, 13 (11), 1066–1071. 10.1038/s41565-018-0232-x.

(9) Chan, A. K.; Chiu, R. W.; Lo, Y. D. Cell-Free Nucleic Acids in Plasma, Serum and Urine: A New Tool in Molecular Diagnosis; 2003; Vol. 40.

(10) Tong, Y. K.; Lo, Y. M. D. Diagnostic Developments Involving Cell-Free (Circulating) Nucleic Acids. Clinica Chimica Acta. January 2006, pp 187–196. 10.1016/j.cccn.2005.05.048.

(11) Hindson, B. J.; Ness, K. D.; Masquelier, D. A.; Belgrader, P.; Heredia, N. J.; Makarewicz, A. J.; Bright, I. J.; Lucero, M. Y.; Hiddessen, A. L.; Legler, T. C.; Kitano, T. K.; Hodel, M. R.; Petersen, J. F.; Wyatt, P. W.; Steenblock, E. R.; Shah, P. H.; Bousse, L. J.; Troup, C. B.; Mellen, J. C.; Wittmann, D. K.; Erndt, N. G.; Cauley, T. H.; Koehler, R. T.; So, A. P.; Dube, S.; Rose, K. A.; Montesclaros, L.; Wang, S.; Stumbo, D. P.; Hodges, S. P.; Romine, S.; Milanovich, F. P.; White, H. E.; Regan, J. F.; Karlin-Neumann, G. A.; Hindson, C. M.; Saxonov, S.; Colston, B. W. High-Throughput Droplet Digital PCR System for Absolute Quantitation of DNA Copy Number. Anal. Chem. 2011, 83 (22), 8604–8610. 10.1021/ac202028g.

(12) Kojabad, A. A.; Farzanehpour, M.; Galeh, H. E. G.; Dorostkar, R.; Jafarpour, A.; Bolandian, M.; Nodooshan, M. M. Droplet Digital PCR of Viral DNA/RNA, Current Progress, Challenges, and Future Perspectives. J. Med. Virol. 2021, 93 (7), 4182–4197. 10.1002/jmv.26846.

(13) Samanta, D.; Ebrahimi, S. B.; Ramani, N.; Mirkin, C. A. Enhancing CRISPR-Cas-Mediated Detection of Nucleic Acid and Non-Nucleic Acid Targets Using Enzyme-Labeled Reporters. J. Am. Chem. Soc. 2022, 144 (36), 16310–16315. 10.1021/jacs.2c07625.

(14) Abudayyeh, O. O.; Gootenberg, J. S.; Kellner, M. J.; Zhang, F. Nucleic Acid Detection of Plant Genes Using CRISPR-Cas13. Cris. J. 2019, 2 (3), 165–171. 10.1089/crispr.2019.0011.

(15) Burkin, K. M.; Ivanov, A. V.; Zherdev, A. V.; Dzantiev, B. B.; Safenkova, I. V. A Critical Study on DNA Probes Attached to Microplate for CRISPR/Cas12 Trans-Cleavage Activity. Biosensors 2023, 13 (8). 10.3390/bios13080824.

(16) Zhu, J.; Kong, J.; Keyser, U. F.; Wang, E. Parallel DNA Circuits by Autocatalytic Strand Displacement and Nanopore Readout. Nanoscale 2022, 14 (41), 15507–15515. 10.1039/d2nr04048d.

(17) Bošković, F.; Zhu, J.; Tivony, R.; Ohmann, A.; Chen, K.; Alawami, M. F.; Đorđević, M.; Ermann, N.; Pereira-Dias, J.; Fairhead, M.; Howarth, M.; Baker, S.; Keyser, U. F. Simultaneous Identification of Viruses and Viral Variants with Programmable DNA Nanobait. Nat. Nanotechnol. 2023, 18 (3), 290–298. 10.1038/s41565-022-01287-x.

(18) Grabenhorst, L.; Pfeiffer, M.; Schinkel, T.; Kümmerlin, M.; Brüggenthies, G. A.; Maglic, J. B.; Selbach, F.; Murr, A. T.; Tinnefeld, P.; Glembockyte, V. Engineering Modular and Tunable Single-Molecule Sensors by Decoupling Sensing from Signal Output. Nat. Nanotechnol. 2024. 10.1038/s41565-024-01804-0.

(19) Yaadav, R.; Trofymchuk, K.; Dass, M.; Behrendt, V.; Hauer, B.; Close, C.; Scheckenbach, M.; Ferrari, G.; Maeurer, L.; Glembockyte, V.; Liedl, T.; Tinnefeld, P. Bringing Attomolar Detection to the Point-of-Care with Nanopatterned DNA Origami Nanoantennas.

(20) Srinivas, N.; Ouldridge, T. E.; Šulc, P.; Schaeffer, J. M.; Yurke, B.; Louis, A. A.; Doye, J. P. K.; Winfree, E. On the Biophysics and Kinetics of Toehold-Mediated DNA Strand Displacement. Nucleic Acids Res. 2013, 41 (22), 10641–10658. 10.1093/nar/gkt801.

(21) Simmel, F. C.; Yurke, B.; Singh, H. R. Principles and Applications of Nucleic Acid Strand Displacement Reactions. Chem. Rev. 2019, 119 (10), 6326–6369. 10.1021/acs.chemrev.8b00580.

(22) Li, S.; Zhu, L.; Lin, S.; Xu, W. Toehold-Mediated Biosensors: Types, Mechanisms and Biosensing Strategies. Biosens. Bioelectron. 2023, 220 (114922). 10.1016/j.bios.2022.114922.

(23) Li, B.; Ellington, A. D.; Chen, X. Rational, Modular Adaptation of Enzyme-Free DNA Circuits to Multiple Detection Methods. Nucleic Acids Res. 2011, 39 (16). 10.1093/nar/gkr504.

(24) Machinek, R. R. F.; Ouldridge, T. E.; Haley, N. E. C.; Bath, J.; Turberfield, A. J. Programmable Energy Landscapes for Kinetic Control of DNA Strand Displacement. Nat. Commun. 2014, 5. 10.1038/ncomms6324.

(25) Bucci, J.; Irmisch, P.; Del Grosso, E.; Seidel, R.; Ricci, F. Orthogonal Enzyme-Driven Timers for DNA Strand Displacement Reactions. J. Am. Chem. Soc. 2022, 144 (43), 19791–19798. 10.1021/jacs.2c06599.

(26) Singh, A.; Patel, G.; Patel, S. S. Twinkle-Catalyzed Toehold-Mediated DNA Strand Displacement Reaction. J. Am. Chem. Soc. 2023. 10.1021/jacs.3c04970.

(27) Fu, J.; Zhang, L.; Long, Y.; Liu, Z.; Meng, G.; Zhao, H.; Su, X.; Shi, S. Multiplexed CRISPR-Based Nucleic Acid Detection Using a Single Cas Protein. Anal. Chem. 2023, 95 (44), 16089–16097. 10.1021/acs.analchem.3c01861.

(28) Gupta, K.; Krieg, E. Y-Switch: A Spring-Loaded Synthetic Gene Switch for Robust DNA/RNA Signal Amplification and Detection. Nucleic Acids Res. 2024. 10.1093/nar/gkae680.

(29) Oishi, M. Comparative Study of DNA Circuit System-Based Proportional and Exponential Amplification Strategies for Enzyme-Free and Rapid Detection of MiRNA at Room Temperature. ACS Omega 2018, 3 (3), 3321–3329. 10.1021/acsomega.7b01866.

(30) Mohammadniaei, M.; Zhang, M.; Ashley, J.; Christensen, U. B.; Friis-Hansen, L. J.; Gregersen, R.; Lisby, J. G.; Benfield, T. L.; Nielsen, F. E.; Henning Rasmussen, J.; Pedersen, E. B.; Olinger, A. C. R.; Kolding, L. T.; Naseri, M.; Zheng, T.; Wang, W.; Gorodkin, J.; Sun, Y. A Non-Enzymatic, Isothermal Strand Displacement and Amplification Assay for Rapid Detection of SARS-CoV-2 RNA. Nat. Commun. 2021, 12 (1). 10.1038/s41467-021-25387-9.

(31) Li, C.; Li, Y.; Xu, X.; Wang, X.; Chen, Y.; Yang, X.; Liu, F.; Li, N. Fast and Quantitative Differentiation of Single-Base Mismatched DNA by Initial Reaction Rate of Catalytic Hairpin Assembly. Biosens. Bioelectron. 2014, 60, 57–63. 10.1016/j.bios.2014.04.007.

(32) Yin, P.; Choi, H. M. T.; Calvert, C. R.; Pierce, N. A. Programming Biomolecular Self-Assembly Pathways. Nature 2008, 451 (7176), 318–322. 10.1038/nature06451.

(33) Dirks, R. M.; Pierce, N. A. Triggered Amplification by Hybridization Chain Reaction. PNAS 2004, 101 (43), 15275–15278. 10.1073/pnas.0407024101.

(34) Choi, H. M. T.; Chang, J. Y.; Trinh, L. A.; Padilla, J. E.; Fraser, S. E.; Pierce, N. A. Programmable in Situ Amplification for Multiplexed Imaging of MRNA Expression. Nat. Biotechnol. 2010, 28 (11), 1208–1212. 10.1038/nbt.1692.

(35) Rösch, E. L.; Sack, R.; Chowdhury, M. S.; Wolgast, F.; Zaborski, M.; Ludwig, F.; Schilling, M.; Viereck, T.; Rand, U.; Lak, A. Amplification- and Enzyme-Free Magnetic Diagnostics Circuit for Whole-Genome Detection of SARS-CoV-2 RNA. ChemBioChem 2024, 25 (16). 10.1002/cbic.202400251.

(36) Zhang, D. Y.; Winfree, E. Control of DNA Strand Displacement Kinetics Using Toehold Exchange. J. Am. Chem. Soc. 2009, 131 (47), 17303–17314. 10.1021/ja906987s.

(37) Long, D.; Shi, P.; Xu, X.; Ren, J.; Chen, Y.; Guo, S.; Wang, X.; Cao, X.; Yang, L.; Tian, Z. Understanding the Relationship between Sequences and Kinetics of DNA Strand Displacements. Nucleic Acids Res. 2024, 52 (16), 9407–9416. 10.1093/nar/gkae652.

(38) Walbrun, A.; Wang, T.; Matthies, M.; Šulc, P.; Simmel, F. C.; Rief, M. Single-Molecule Force Spectroscopy of Toehold-Mediated Strand Displacement. Nat. Commun. 2024, 15 (1), 1–15. 10.1038/s41467-024-51813-9.

(39) Irmisch, P.; Ouldridge, T. E.; Seidel, R. Modeling DNA-Strand Displacement Reactions in the Presence of Base-Pair Mismatches. J. Am. Chem. Soc. 2020, 142 (26), 11451–11463. 10.1021/jacs.0c03105.

(40) Haley, N. E. C.; Ouldridge, T. E.; Mullor Ruiz, I.; Geraldini, A.; Louis, A. A.; Bath, J.; Turberfield, A. J. Design of Hidden Thermodynamic Driving for Non-Equilibrium Systems via Mismatch Elimination during DNA Strand Displacement. Nat. Commun. 2020, 11 (1). 10.1038/s41467-020-16353-y.

(41) Zhang, X. L.; Yang, Z. H.; Chang, Y. Y.; Liu, D.; Li, Y. R.; Chai, Y. Q.; Zhuo, Y.; Yuan, R. Programmable Mismatch-Fueled High-Efficiency DNA Signal Converter. Chem. Sci. 2020, 11 (1), 148–153. 10.1039/c9sc05084a.

(42) Oliveira, L. M.; Long, A. S.; Brown, T.; Fox, K. R.; Weber, G. Melting Temperature Measurement and Mesoscopic Evaluation of Single, Double and Triple DNA Mismatches. Chem. Sci. 2020, 11 (31), 8273–8287. 10.1039/d0sc01700k.

(43) Broadwater, D. W. B.; Kim, H. D. The Effect of Basepair Mismatch on DNA Strand Displacement. Biophys. J. 2016, 110 (7), 1476–1484. 10.1016/j.bpj.2016.02.027.

(44) Qiu, G.; Gai, Z.; Tao, Y.; Schmitt, J.; Kullak-Ublick, G. A.; Wang, J. Dual-Functional Plasmonic Photothermal Biosensors for Highly Accurate Severe Acute Respiratory Syndrome Coronavirus 2 Detection. ACS Nano 2020, 14 (5), 5268–5277. 10.1021/acsnano.0c02439.

(45) Taton, T. A.; Mirkin, C. A.; Letsinger, R. L. Scanometric DNA Array Detection with Nanoparticle Probes; 2000; Vol. 289. 10.1126/science.289.5485.1757.

(46) Zhang, X.; Reeves, D. B.; Perreard, I. M.; Kett, W. C.; Griswold, K. E.; Gimi, B.; Weaver, J. B. Molecular Sensing with Magnetic Nanoparticles Using Magnetic Spectroscopy of Nanoparticle Brownian Motion. Biosens. Bioelectron. 2013, 50, 441–446. 10.1016/j.bios.2013.06.049.

(47) Alafeef, M.; Dighe, K.; Moitra, P.; Pan, D. Rapid, Ultrasensitive, and Quantitative Detection of SARS-CoV-2 Using Antisense Oligonucleotides Directed Electrochemical Biosensor Chip. ACS Nano 2020, 14 (12), 17028–17045. 10.1021/acsnano.0c06392.

(48) Sanz-De Diego, E.; Aires, A.; Palacios-Alonso, P.; Cabrera, D.; Silvestri, N.; Vequi-Suplicy, C. C.; Artés-Ibáñez, E. J.; Requejo-Isidro, J.; Delgado-Buscalioni, R.; Pellegrino, T.; Cortajarena, A. L.; Terán, F. J.; Cortajarena, A. L. Multiparametric Modulation of Magnetic Transduction for Biomolecular Sensing in Liquids. Nanoscale 2024, 16 (8), 4082–4094. 10.1039/d3nr06489a.

(49) Lak, A.; Wang, Y.; Kolbeck, P. J.; Pauer, C.; Chowdhury, M. S.; Cassani, M.; Ludwig, F.; Viereck, T.; Selbach, F.; Tinnefeld, P.; Schilling, M.; Liedl, T.; Tavacoli, J.; Lipfert, J. Cooperative Dynamics of DNA-Grafted Magnetic Nanoparticles Optimize Magnetic Biosensing and Coupling to DNA Origami. Nanoscale 2024, 16 (15), 7678–7689. 10.1039/d3nr06253h.

(50) Haun, J. B.; Yoon, T. J.; Lee, H.; Weissleder, R. Magnetic Nanoparticle Biosensors. Wiley Interdiscip. Rev. Nanomedicine Nanobiotechnology 2010, 2 (3), 291–304. 10.1002/wnan.84.

(51) Chung, S. H.; Hoffmann, A.; Bader, S. D.; Liu, C.; Kay, B.; Makowski, L.; Chen, L. Biological Sensors Based on Brownian Relaxation of Magnetic Nanoparticles. Appl. Phys. Lett. 2004, 85 (14), 2971–2973. 10.1063/1.1801687.

(52) Krishnan, K. M. Biomedical Nanomagnetics: A Spin through Possibilities in Imaging, Diagnostics, and Therapy. IEEE Trans. Magn. 2010, 46 (7), 2523–2558. 10.1109/TMAG.2010.2046907.

(53) Wu, K.; Liu, J.; Chugh, V. K.; Liang, S.; Saha, R.; Krishna, V. D.; Cheeran, M. C. J.; Wang, J. P. Magnetic Nanoparticles and Magnetic Particle Spectroscopy-Based Bioassays: A 15 Year Recap. Nano Futur. 2022, 6 (2), 022001. 10.1088/2399-1984/ac5cd1.

(54) Orlov, A. V.; Znoyko, S. L.; Cherkasov, V. R.; Nikitin, M. P.; Nikitin, P. I. Multiplex Biosensing Based on Highly Sensitive Magnetic Nanolabel Quantification: Rapid Detection of Botulinum Neurotoxins A, B, and e in Liquids. Anal. Chem. 2016, 88 (21), 10419–10426. 10.1021/acs.analchem.6b02066.

(55) Krause, H. J.; Engelmann, U. M. Fundamentals and Applications of Dual-Frequency Magnetic Particle Spectroscopy: Review for Biomedicine and Materials Characterization. Adv. Sci. 2025, 2416838. 10.1002/advs.202416838.

(56) Crowell, E.; Geng, L. Reduction of Multiexponential Background in Fluorescence with Phase-Sensitive Detection. Appl. Spectrosc. 2001, 55 (12), 1709–1716. 10.1366/0003702011954062.

(57) Pandey, V.; Pandey, T. Biophysical Significance of Fluorescence Spectroscopy in Deciphering Nucleic Acid Dynamics: From Fundamental to Recent Advancements. Biophys. Chem. 2025, 316 (October 2024). 10.1016/j.bpc.2024.107345.

(58) Issadore, D.; Park, Y. I.; Shao, H.; Min, C.; Lee, K.; Liong, M.; Weissleder, R.; Lee, H. Magnetic Sensing Technology for Molecular Analyses. Lab Chip 2014, 14 (14), 2385–2397. 10.1039/c4lc00314d.

(59) Lin, G.; Makarov, D.; Schmidt, O. G. Magnetic Sensing Platform Technologies for Biomedical Applications. Lab Chip 2017, 17 (11), 1884–1912. 10.1039/c7lc00026j.

(60) Rauwerdink, A. M.; Weaver, J. B. Measurement of Molecular Binding Using the Brownian Motion of Magnetic Nanoparticle Probes. Appl. Phys. Lett. 2010, 96 (3). 10.1063/1.3291063.

(61) Tian, B.; Han, Y.; Wetterskog, E.; Donolato, M.; Hansen, M. F.; Svedlindh, P.; Strömberg, M. MicroRNA Detection through DNAzyme-Mediated Disintegration of Magnetic Nanoparticle Assemblies. ACS Sensors 2018, 3 (9), 1884–1891. 10.1021/acssensors.8b00850.

(62) Kahmann, T.; Wolgast, F. T.; Viereck, T.; Schilling, M.; Ludwig, F. Improvements of Magnetic Nanoparticle Assays for SARS-CoV-2 Detection Using a Mimic Virus Approach. Sens. Bio-Sensing Res. 2024, 44. 10.1016/j.sbsr.2024.100654.

(63) Rösch, E. L.; Zhong, J.; Lak, A.; Liu, Z.; Etzkorn, M.; Schilling, M.; Ludwig, F.; Viereck, T.; Lalkens, B. Point-of-Need Detection of Pathogen-Specific Nucleic Acid Targets Using Magnetic Particle Spectroscopy. Biosens. Bioelectron. 2021, 192. 10.1016/j.bios.2021.113536.

(64) Wu, K.; Chugh, V. K.; di Girolamo, A.; Liu, J.; Saha, R.; Su, D.; Krishna, V. D.; Nair, A.; Davies, W.; Wang, Y. A.; Cheeran, M. C. J.; Wang, J. P. A Portable Magnetic Particle Spectrometer for Future Rapid and Wash-Free Bioassays. ACS Appl. Mater. Interfaces 2021, 13 (7), 7966–7976. 10.1021/acsami.0c21040.

(65) Wolgast, F. T.; Kahmann, T.; Janssen, K. J.; Yoshida, T.; Zhong, J.; Schilling, M.; Ludwig, F.; Viereck, T. Low-Cost Magnetic Particle Spectroscopy Hardware for Low-Viral-Load Immunoassays. IEEE Trans. Instrum. Meas. 2024. 10.1109/TIM.2024.3449985.

(66) Li, T.; Meng, F.; Fang, Y.; Luo, Y.; He, Y.; Dong, Z.; Tian, B. Multienzymatic Disintegration of DNA-Scaffolded Magnetic Nanoparticle Assembly for Malarial Mitochondrial DNA Detection. Biosens. Bioelectron. 2024, 246. 10.1016/j.bios.2023.115910.

(67) Huang, Z.; Li, J.; Zhong, H.; Tian, B. Nucleic Acid Amplification Strategies for Volume-Amplified Magnetic Nanoparticle Detection Assay. Front. Bioeng. Biotechnol. 2022, 10. 10.3389/fbioe.2022.939807.

(68) Martín, D. S.; Oropesa-Nuñez, R.; de la Torre, T. Z. G. Rolling Circle Amplification on a Bead: Improving the Detection Time for a Magnetic Bioassay. ACS Omega 2023, 8 (4), 4391–4397. 10.1021/acsomega.2c07992.

(69) Poppleton, E.; Matthies, M.; Mandal, D.; Romano, F.; Šulc, P.; Rovigatti, L. OxDNA: Coarse-Grained Simulations of Nucleic Acids Made Simple. J. Open Source Softw. 2023, 8 (81), 4693. 10.21105/joss.04693.

(70) Snodin, B. E. K.; Randisi, F.; Mosayebi, M.; Šulc, P.; Schreck, J. S.; Romano, F.; Ouldridge, T. E.; Tsukanov, R.; Nir, E.; Louis, A. A.; others. Introducing Improved Structural Properties and Salt Dependence into a Coarse-Grained Model of DNA. J. Chem. Phys. 2015, 142 (23).

(71) Oishi, M. Enzyme-Free and Isothermal Detection of MicroRNA Based on Click-Chemical Ligation-Assisted Hybridization Coupled with Hybridization Chain Reaction Signal Amplification. Anal. Bioanal. Chem. 2015, 407 (14), 4165–4172. 10.1007/s00216-015-8629-y.

(72) Cisse, I. I.; Kim, H.; Ha, T. A Rule of Seven in Watson-Crick Base-Pairing of Mismatched Sequences. Nat. Struct. Mol. Biol. 2012, 19 (6), 623–627. 10.1038/nsmb.2294.

(73) Ludwig, F.; Guillaume, A.; Schilling, M.; Frickel, N.; Schmidt, A. M. Determination of Core and Hydrodynamic Size Distributions of CoFe 2O4 Nanoparticle Suspensions Using Ac Susceptibility Measurements. J. Appl. Phys. 2010, 108 (3). 10.1063/1.3463350.

(74) Ludwig, F.; Balceris, C.; Jonasson, C.; Johansson, C. Analysis of AC Susceptibility Spectra for the Characterization of Magnetic Nanoparticles. IEEE Trans. Magn. 2017, 53 (11), 11–14. 10.1109/TMAG.2017.2693420.

(75) Rossetti, G.; Dans, P. D.; Gomez-Pinto, I.; Ivani, I.; Gonzalez, C.; Orozco, M. The Structural Impact of DNA Mismatches. Nucleic Acids Res. 2015, 43 (8), 4309–4321. 10.1093/nar/gkv254.

(76) Li, C.; Li, Y.; Chen, Y.; Lin, R.; Li, T.; Liu, F.; Li, N. Modulating the DNA Strand-Displacement Kinetics with the One-Sided Remote Toehold Design for Differentiation of Single-Base Mismatched DNA. RSC Adv. 2016, 6 (78), 74913–74916. 10.1039/c6ra17006d.

(77) Genot, A. J.; Zhang, D. Y.; Bath, J.; Turberfield, A. J. Remote Toehold: A Mechanism for Flexible Control of DNA Hybridization Kinetics. J. Am. Chem. Soc. 2011, 133 (7), 2177–2182. 10.1021/ja1073239.

(78) Seeman, N. C. The Design of DNA Sequences for Branched Systems. In Structural DNA Nanotechnology; 2015; pp 11–27. 10.1017/cbo9781139015516.003.

(79) Yang, X.; Tang, Y.; Traynor, S. M.; Li, F. Regulation of DNA Strand Displacement Using an Allosteric DNA Toehold. J. Am. Chem. Soc. 2016, 138 (42), 14076–14082. 10.1021/jacs.6b08794.

(80) SantaLucia, J.; Hicks, D. The Thermodynamics of DNA Structural Motifs. Annu. Rev. Biophys. Biomol. Struct. 2004, 33, 415–440. 10.1146/annurev.biophys.32.110601.141800.

(81) Owczarzy, R.; You, Y.; Groth, C. L.; Tataurov, A. V. Stability and Mismatch Discrimination of Locked Nucleic Acid-DNA Duplexes. Biochemistry 2011, 50 (43), 9352–9367.

(82) Olson, X.; Kotani, S.; Yurke, B.; Graugnard, E.; Hughes, W. L. Kinetics of DNA Strand Displacement Systems with Locked Nucleic Acids. J. Phys. Chem. B 2017, 121 (12), 2594–2602. 10.1021/acs.jpcb.7b01198.

