## Supplementary material for "Mismatch-Assisted Toehold Exchange Cascades for Magnetic Nanoparticle-Based Nucleic Acid Diagnostics": SI

### Experimental procedures

#### Materials

BNF80 (micromod particle technology GmbH) MNPs, which are bionized NanoFerrite crystallites in a starch shell functionalized with Streptavidin, were used in this work. The DNA sequences were purchased from eurofins genomics GmbH. DNA sequences containing a locked nucleic acid (LNA) were purchased from Integrated DNA Technologies Inc. Lyophilized DNA was rehydrated in custom buffer at a starting concentration of 100 µM. The custom sample buffer contains 150 mM NaCl (Merck KGaA), 20 mM Tris-HCl, 1 mM EDTA and 1 % Triton-X (Carl Roth GmbH + Co. KG) in MilliQ water. NaOH and HCl are used to adjust the pH value to 7.4. $MgCl_2$ (Carl Roth GmbH + Co. KG) is dissolved in the sample buffer at a concentration of 1 µM for further dilution.

#### Synthesis procedures

Clusters were formed by cross-linking functionalized MNPs, labelled separately with two distinct Label DNA sequences, in equal ratio with complementary DNA (substrate). The steps from the bare building blocks up to the measurement are described in the following sections.

##### DNA design choices

The target (T) sequence (table S1) used in this work was adapted from Mohammadniaei et al. [1] and consists of domains $TH_T$ and BM (Figure 1b). The sequence of domain a ($L_{Stay}$) was adopted from a previous magnetic assay design after analyzing the intended hybridization reactions with NUPACK[1]. The label DNA was designed with a poly adenine (A) spacer (10A), as it improves hybridization [2]. Strands $L_{Stay}$ and $L_{DSD}$ are modified with Biotin-TEG to allow a sufficient distance from the MNP surface to enhance the surface functionalization[2,3]. Domain b* was elongated from 7 to 8 nt to investigate different lengths α of $TH_T$, counted from the 5′ end of domain b*, from 7 to 5 nt, while reducing potential stacking interactions of $TH_T$ to the double-stranded domain a of $L_{Stay}$ [3]. A singular A nucleotide was added to avoid hairpin formation in b*, which would otherwise interfere significantly with TH initiation. In this work, a target TH length of $\alpha = 7$ nt is utilized, unless stated differently. Sequences of mismatched substrates Sx and their corresponding complementary fuels Fx are listed in table S2.

---

[1] https://www.nupack.org/
[2] https://eu.idtdna.com/pages/education/decoded/article/which-biotin-modification-to-use-
[3] https://eurofinsgenomics.eu/en/dna-rna-oligonucleotides/custom-dna-rna-oligos/custom-dna-oligos/

**Table S1. DNA sequences used in this work.** Domains separated by "-", last bases of T for β = 0 bp in brackets.

| Strand identity | Domain | Nucleic acid sequence (5´ -> 3´) |
|---|---|---|
| $L_{DSD}$ | c | ACC CGC AAT CCT GCT AAC (AAAAA AAAAA) BioTEG |
| $L_{Stay}$ | a | BioTEG (AAAAA AAAAA) TA ATT GTA TGT GTG TAG C |
| $S_0$ | c* - b* - a* | GTT AGC AGG ATT GCG GGT - GCC AAT GA - GCT ACA CAC ATA CAA TTA |
| $F_0$ | BM - $TH_F$ | ACC CGC AAT CCT - GCT AAC |
| T (α = 7, β = 6 bp) | $TH_T$ - BM(-$TH_F$) | C ATT GGC - ACC CGC AAT CCT (- GCT AAC) |

**Table S2. Sequence and free energy of DNA used in studies on mismatch positions.** Sequences of substrate DNA Sx including an underlined mismatched bp at position x between Sx and $L_{DSD}$, as well as sequences of fuel DNA Fx, complementary to Sx. Free energy Δ*G* [kcal/mol] of DNA structures calculated via NUPACK simulations (150 mM $Na^+$, 0 mM $Mg^{2+}$, 5 nM oligo concentration, 25 °C, all stacking).

| Strand identity | Sequence (5´ -> 3´) | $L_{DSD}$ + Sx + $L_{Stay}$ | $L_{Stay}$ + Sx + T (α = 7, β = 6 bp) | $L_{Stay}$ + Sx + Fx |
|---|---|---|---|---|
| S0<br><br>F0 | GTT AGC AGG ATT GCG GGT-GCC AAT GA-G CTA CAC ACA TAC AAT TA<br><br>ACC CGC AAT CCT GCT AAC | -51.94 | -53.55 | -50.83 |
| S2<br><br>F2 | G**G**T AGC AGG ATT GCG GGT-GCC AAT GA-G CTA CAC ACA TAC AAT TA<br><br>ACC CGC AAT CCT GCT A**C**C | -49.54 | -53.55 | -51.64 |
| S3<br><br>F3 | GT**G** AGC AGG ATT GCG GGT-GCC AAT GA-G CTA CAC ACA TAC AAT TA<br><br>ACC CGC AAT CCT GCT **C**AC | -48.79 | -53.55 | -52.01 |
| S4<br><br>F4 | GTT **C**GC AGG ATT GCG GGT-GCC AAT GA-G CTA CAC ACA TAC AAT TA<br><br>ACC CGC AAT CCT GC**G** AAC | -48.48 | -53.55 | -52.52 |
| S5<br><br>F5 | GTT A**A**C AGG ATT GCG GGT-GCC AAT GA-G CTA CAC ACA TAC AAT TA<br><br>ACC CGC AAT CCT G**T**T AAC | -46.72 | -53.55 | -49.74 |
| S6<br><br>F6 | GTT AG**A** AGG ATT GCG GGT-GCC AAT GA-G CTA CAC ACA TAC AAT TA<br><br>ACC CGC AAT CCT **T**CT AAC | -47.33 | -53.61 | -49.41 |

#### Functionalization of MNPs with label DNA

BNF80 MNPs (1.6 nM) and label DNA (104.8 nM) were mixed in sample buffer to a total volume of 281.4 µl. This was done once for each type of label DNA ($L_{Stay}$ and $L_{DSD}$), see table S2 for the sequences. Each ingredient was carefully vortexed before pipetting to ensure homogeneous preparation. After combining the elements, the tubes were vortexed and sonicated for 10 s to break potential MNP agglomerations. We then incubated the samples in a thermomixer for 4 h at 25 °C, with a 10-min cycle of 650 rpm for 5 min and break intervals of 5 min. The DNA-labelled MNPs were then washed to remove any unbound label DNA by centrifugation at 3000 rcf and 21 °C for 25 min. Next, we discarded 260 µl of the supernatant. The same volume of fresh sample buffer is added to the particle pellet, which was then vortexed thoroughly to fully redisperse the MNPs. The washing step was repeated one more time. The added volume was reduced to reach a total volume of 237 µl to compensate for the loss of MNPs during the washing process of 16 %, which was determined via inductively coupled plasma optical emission spectroscopy (ICP-OES) [4]. The collected MNPs were fully dispersed by thorough vortexing and sonication for 10 s.

**Formation of magnetic clusters**

In a typical cluster synthesis procedure, 118 µl of $L_{Stay}$-labelled MNPs, 118 µl of $L_{DSD}$-labelled MNPs and 37.8 µl substrate (0.1 µM) were added to 198.2 µl sample buffer, yielding a total volume of 472 µl with 0.8 nM BNF80 and 8 nM substrate. The clusters incubate in a thermomixer for 18 h at 25 °C, with a 10-min cycle of 650 rpm for 5 min and break intervals of 5 min. The resulting magnetic clusters were washed by two rounds of centrifugation at 1000 rcf and 4 °C for 5 min to remove any unbound substrates. After each round, 450 µl supernatant was pipetted out. We adjusted the sample volume to 397 µl after the final washing step to compensate for 16 % particle loss. The clusters were then vortexed vigorously to break unspecific MNP agglomerations. Purified magnetic clusters were stored at 4 °C prior to further use.

While the cluster formation is reproducible with only small deviations due to MNP agglomerations as a result of centrifugation, data within one experiment were sourced from one preparation batch to ensure optimal comparability. One set of DNA-labelled MNPs yields two tubes of clusters, which each allow the preparation of five samples for declustering-based assays over time via an ACS setup or ten samples to be measured with our custom MPS setup. Experiments of bigger sample number are prepared by combining the required volume of functionalized MNPs and clusters after the respective purification step.

#### Principles of magnetic assays

Solution-based magnetic detection of nucleic acid sequences is based on the dynamic response of Brownian relaxing MNPs to alternating magnetic fields. The internal magnetic moment of these MNPs is blocked along a specific crystalline axis of particles. To align their magnetic moment to external magnetic fields, these MNPs must physically rotate, a process called Brownian relaxation. The relaxation time constant $\tau_B$ of the Brownian relaxation processes is given by

$$\tau_B = \frac{3\eta V_H}{k_B T} \quad (1)$$

with $\eta$ the viscosity of the sample, $V_H$ hydrodynamic volume of the MNPs, $k_B$ Boltzmann constant and $T$ temperature. When a biomarker is attached to a MNP, it's $V_H$ increases, which increases its $\tau_B$ and can be detected via magnetic measurements. The increase in $V_H$ is prone to false-positive results as a result of nonspecific agglomeration of MNPs due to electrostatic and magnetic interactions. Therefore, we have designed a completely different detection concept, in which the disassembly of magnetic clusters made of MNPs, that are connected to each other via complementary substrate DNA, is monitored. In the presence of a specific target nucleic acid sequence, the clusters are broken up into single MNPs via TMSD, decreasing the hydrodynamic size in a highly specific manner.

The properties of the magnetic excitation field are crucial for the magnetic read-out and the sensitivity of the magnetic assays [5]. Therefore, amplitude $\hat{H}$ and frequency $f$ of the sinusoidal magnetic field were tailored specifically for BNF80 MNPs [6].

**Kinetic studies using complex ac susceptometry**

In a complex alternating current susceptometry (ACS) measurement both the real ($\chi'$) and imaginary ($\chi''$) parts of ACS are measured. For our kinetics studies, we only look at the imaginary part $\chi''$, which is expressed by

$$\chi''(\omega) = \frac{\chi_0 \omega \tau_B}{1+(\omega \tau_B)^2} \quad (2)$$

with the static initial susceptibility $\chi_0$ and the excitation frequency $\omega$. The static susceptibility

$$\chi_0 = \frac{\mu_0 n \mu_{\mathrm{eff}}^2}{3 k_B T} \quad (3)$$

is determined by the vacuum magnetic permeability $\mu_0$, number of MNPs $n$ and $\mu_{\mathrm{eff}}$ effective magnetic moment of the MNPs [7] [Ludwig2010]. For a system of MNPs in equilibrium, $\chi''(\omega)$ peaks at the characteristic frequency $\omega=1/\tau_B$. The peak frequency and amplitude depend directly on the particle hydrodynamic size [8], see Figure S1a for the $\chi''$ distribution of magnetic clusters of varying sizes. The ACS measurements in this work were executed with a custom setup at an excitation field of $f$=120 Hz and $\hat{H}$=0.5 mT/$\mu_0$, recording a data-point every 23.75 s. The field settings were chosen to capture the highest possible signal variation during the disassembly of the magnetic clusters (Figure S1b). An increase of $\chi''$ at 120 Hz is caused by the release of MNPs from clusters and the consequential decrease in overall hydrodynamic size of the magnetic clusters (Figure S1a). No background measurement is needed due to the design of the setup. The measured data of different clusters are plotted as the relative change $\Delta\chi''$ normalized to the first data point $\chi''_0 = \chi''(0 \text{ min})$, which is calculated by

$$\frac{\Delta\chi''}{\chi''_0} = \frac{\chi''(t) - \chi''_0}{\chi''_0} \tag{4}$$

to present the overall signal change and eliminate the influence of differences between different cluster batches on the measured value $\chi''$.

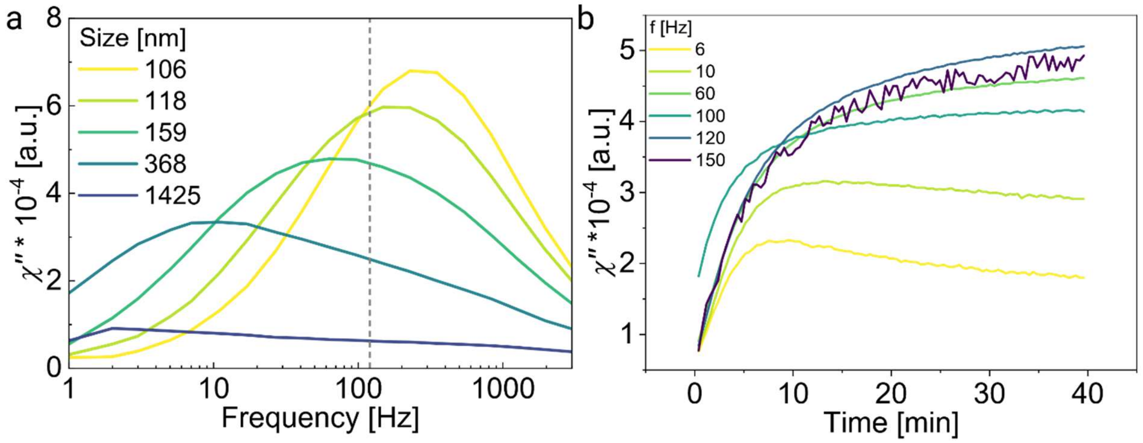

**Figure S1. Supporting ACS measurements.** (a) Exemplary spectra of the ACS imaginary part of magnetic clusters, formed from BNF80 MNPs by adding different concentrations of substrate, which increases the hydrodynamic size. The $\chi''$ peak shifts to lower frequency and decreases in amplitude with size $V_H$ of the magnetic clusters. A dashed line marks the frequency of 120 Hz, at which the kinetic measurements in this study were executed, demonstrating the enormous increase in $\chi''$ during declustering processes. Hydrodynamic sizes were determined via Dynamic Light Scattering (Malvern Zetasizer). (b) $\chi''$ measured for different excitation frequencies $f = \frac{\omega}{2\pi}$ at a field strength of 0.5 mT, for an excess of target to substrate, with full displacement of the incumbent DNA strands.

The samples for the ACS measurement required 150 µl volume. The pipetting order of the ingredients was: custom buffer, magnesium (1 M), 75 µl clusters, fuel DNA and target DNA. The measured samples contained 0.4 nM MNPs, 4 nM substrate DNA and 100 mM $Mg^{2+}$ (unless stated differently), as well as varying concentrations of DNA inputs, which are specified in the respective result chapter. Elements which are not required in the experiment were skipped. The additives were vortexed before they were added. For the ACS measurements over time, the sample was vortexed briefly after the last ingredient was added. We then pipetted 140 µl into a measurement vial, transferred it to our custom ACS setup and started the measurement. We acknowledge the presence of a short time delay, which is inevitable to avoid potential magnetic contamination of the coil setup surrounding the sample holder. The sample is moved vertically within the magnetic coils, therefore adding the last DNA element via pipette-mixing is not feasible.

**Snapshot measurements via MPS for LoD determination**

Our custom MPS system operates at a frequency of 590 Hz and $\widehat{H}$=15 mT/$\mu_0$ [6]. The system calculates the amplitudes of the uneven higher harmonics of the excitation frequency that are characteristic for colloidal MNPs with specific properties [9]. The amplitudes increase with decreasing $V_H$ and increasing particle concentration. To rule out errors that may be caused by variations in the MNP concentration, the ratio of the 5th ($H_5$) to the 3rd ($H_3$) harmonics amplitude

$$HR_{53} = H_5 / H_3 \tag{5}$$

was calculated and used to evaluate the assay. The $HR_{53}$ increases with decreasing $V_H$ and can therefore be utilized to detect declustering processes [10]. Prior to each sample measurement, a background sample was measured and subtracted automatically. Each sample was measured five times, and the average and standard errors were calculated. The relative change $\Delta HR_{53}$ was calculated via

$$\Delta HR_{53} = \frac{HR_{53,\text{sample}} - HR_{53,\text{control}}}{HR_{53,\text{control}}} \tag{6}$$

to normalize the results of $HR_{53,\text{sample}}$ to a control sample ($HR_{53,\text{control}}$), which was prepared without target DNA as reference.

The samples for the MPS measurements were prepared with 75 µl sample volume. The pipetting order of the ingredients was: custom buffer, magnesium (1 M), 37.5 µl clusters, fuel DNA and target DNA. The measured samples contained 0.4 nM MNPs, 4 nM substrate DNA and 100 mM $Mg^{2+}$ (unless stated differently), as well as varying concentrations of DNA inputs, which are specified in the respective result chapter. Elements which are not required in the experiment were skipped. The additives were vortexed before they were added. Once the last ingredient is added to the sample volume, the sample is vortexed and transferred to a thermomixer. The samples are incubated at 25 °C, with 10-min cycles: 2.5 min at 1400 rpm and 7.5 min break intervals. We use a higher rpm number to avoid sedimentation of clusters over longer periods of time. After thorough vortexing we then transfer 60 µl of the sample volume to a glass vial for the MPS measurement.

#### oxDNA simulations

**Forward Flux Sampling**

We used Forward Flux Sampling (FFS) [11], using a coarse-grained model oxDNA [12,13] to estimate the rate of the dissociation of the DNA strand bound to the substrate. The FFS simulation tool follows the typical setup procedure, used in prior works where we simulated a kinetic process in DNA nanotechnology using oxDNA [14,15].

We defined an order parameter for the FFS algorithm as the number of hydrogen bonds. In order to ensure dissociation, our order parameter included all possible hydrogen bonds between the six complementary bases of the substrate and the entire label strand. We also defined an order parameter for the minimum distance between any two of the bases included in the bonding order parameter.

We partitioned the dissociation reaction with four interfaces $\lambda_1$, $\lambda_2$, $\lambda_3$ and $\lambda_4$. We defined $\lambda_1$ as the point where the system had two fewer hydrogen bonds (as defined in our order parameter) compared to the initial state, $\lambda_2$ as the state where the system had three hydrogen bonds, $\lambda_3$ as the state where the system had one hydrogen bond, and $\lambda_4$ was defined as the state where the minimum distance between nucleotides that could form a bond was greater than 3.4 nanometers.

We performed the FFS simulations for different dissociating strands with different position of mismatch between $L_{DSD}$ and the substrate (second, third, fourth, fifth or sixth base, or no mismatch). For each studied case, we performed three independent replicas of the FFS simulations to quantify the error of the estimated rate of dissociation, as calculated from the FFS simulations. All FFS simulations were run at 25 °C.

**Free Energy Profile estimated from VMMC simulations**

We used oxDNA to perform Virtual Move Monte Carlo (VMMC) free energy profiling of the dissociation of the label from the substrate, which is a procedure to estimate from the simulations the probability of observing a particular number of base pairs in the system at equilibrium. The free energy $F(x)/k_B T$ in the profile is equal to (up to a constant) to the logarithm of probability of the system having $x$ bases. All our simulations were performed at a temperature of 25 °C. To ensure that the system visited all states which we wanted to profile, we used umbrella sampling to ensure that simulations samples states which would otherwise be less free-energetically favorable.

We performed fifteen independent replicas of each mismatch case (no mismatch, mismatch at position 2, mismatch at position 3, mismatch at position 4, mismatch at position 5, mismatch at position 6). Each simulation produced a histogram of the number of snapshots taken of the system in each state. These numbers were then unbiased to remove the effects of the weights used in the VMMC simulation. We then calculated the proportion of time spent in each state by dividing the unbiased weights by the total number of snapshots, and from there computed the free energy (in units of $k_B T$) of each state was by taking the negative logarithm of the proportion of snapshots in the state. Our set up and evaluations of the umbrella sampling VMMC simulations with oxDNA follows the procedures used previously with oxDNA for DNA nanotechnology systems [16]. To be able to sample relative free-energy difference, we ran the simulation at strand concentrations corresponding to $3.34 \times 10^{-4}$ M concentration, much larger than the strand concentration used in experiments. However, the relative difference between the stability of the bound state between different mismatch positions in the simulation is independent of the concentration.

In order to compute the error associated with the free energies, we divided the fifteen replicas up into three groups of five simulations. We computed the average of each of the three groups, as well as the energy of the free energy as the average difference between pairs of subgroups. When plotting the free energies, we shifted the curves along the Y-axis so that the free energies are all shown relative to zero free when the system has zero hydrogen bonds.

The resulting free-energy profiles are shown in Figure 2d. We set up our simulations and performed these calculations with the help of the ipy_oxDNA python library [17].

### Supplementary Data

#### Limited declustering for partial displacement of $L_{DSD}$

In a first attempt to employ TH-exchange in magnetic cascades, we studied the influence of the length of the spontaneous dissociation domain $TH_F$ (β) on the declustering behavior. We measured $\chi''$ over 45 min and $HR_{53}$ after 5 h of incubation (Figure S2).

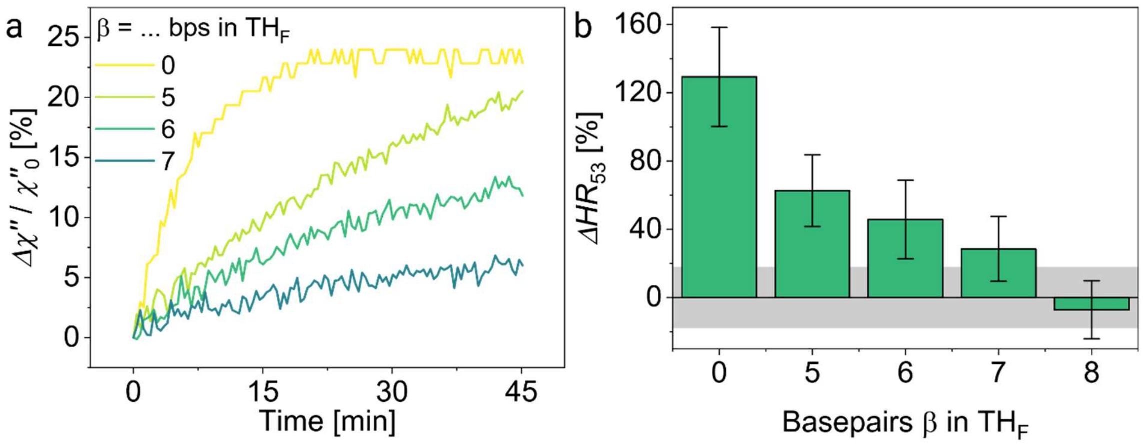

**Figure S2. Limited declustering for partial displacement of $L_{DSD}$ in the absence of a mismatched bp.** Experiments were executed on clusters with S0 with varying lengths β of $TH_F$, modulating the number of bps to dissociate spontaneously for a MNP to detach and a signal change to occur (α = 7 bp). a) Normalized $\chi''$ in the presence of 1 nM T, calculated via equation 4 (100 mM $Mg^{2+}$, 4 nM S0). b) $\Delta HR_{53}$ calculated via equation 6, measured after 5 h incubation at 25 °C (10 mM $Mg^{2+}$, 3 nM T, 3 nM S0); grey marked area 1σ of control sample.

The steepest increase of the samples, as well as saturation in $\Delta\chi''$ (Figure S2a) can be observed when the full length of $L_{DSD}$ is displaced (β = 0 bp). For β = 5-7 bp, spontaneous dissociation is required for MNPs to be released and a change in $\chi''$ to be monitored. The measured $\chi''$ values of these samples increase significantly slower and show lower overall signal change, implying that the declustering occurs incompletely and at a decelerated pace. The same effects were observed via MPS measurements after 5 h of incubation (Figure S2b), with a limited signal change $\Delta HR_{53}$ for all samples requiring spontaneous dissociation. For β = 8 bp, no relevant signal change was observed at all, since the ΔHR53 of the sample did not exceed the 1σ level of the control sample.

The shorter length of β = 5 bp would offer faster spontaneous dissociation than β = 6 bp for a complementary duplex, but the length of $TH_F$ would be too short for the recycling to initiate well. For a spontaneous dissociation of β = 7 bp, little to no declustering was observed, matching findings in literature [18]. Two main effects come together to cause this behavior: first, a longer spontaneous dissociation domain is more stable and detaches slower. Second, the equilibrium of the dsDNA product is less energetically favorable for a longer spontaneous dissociation domain, since less base-pairs are gained in the TMSD reaction. We continued our study with a spontaneous dissociation of β = 6 bp, since this length promises viable TH initiation for the recycling TMSD reaction and may be enhanced via introduction of a mismatched base-pair.

#### Mismatch position dependent rates pre-averaging

Rates for declustering in Figure 2c were averaged two experimental series (Figure S3).

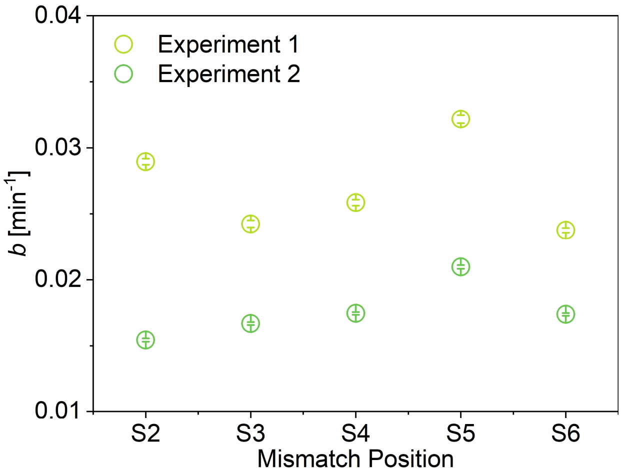

**Figure S3. Kinetic rates for different mismatch positions Sx.** Experimentally derived parameter *b*.

While the experimentally derived rates *b* differ greatly between the two independent experiment series, leading to big error bars in Figure 2c, the trends between the MM positions are consistent. The differences between the series can be explained through variations within the used MNP batches, in terms of their particle size, which in turn influences cluster size and the magnetic signal. Potential fluctuations in room temperature due to a time span of 3 months between these two series may also partially explain deviations seen between the two series.

**Table S3. Fit parameters for the comparison of spontaneous dissociation of 6 bps in the presence of different mismatch positions x.** An exponential fit of the normalized data, shown averaged in Figure 2b, was executed via equation 1, listed are the estimated total magnetic signal gain *A* [%] and the saturation rate *b* [$min^{-1}$] of the individual data sets.

| Substrate identity | *A* [%] | | *b* [$min^{-1}$] | |
|---|---|---|---|---|
| | Set 1 | Set 2 | Set 1 | Set 2 |
| S0 | 13.2 | 28.7 | 0.06646 | 0.02934 |
| S2 | 73.4 | 88.4 | 0.02897 | 0.01543 |
| S3 | 92.7 | 107.2 | 0.02422 | 0.01667 |
| S4 | 84.1 | 122.2 | 0.02584 | 0.01743 |
| S5 | 93.7 | 106.1 | 0.03218 | 0.02096 |
| S6 | 142.9 | 152.9 | 0.02374 | 0.01736 |

#### Declustering in the presence of a blocker strand

We observed increasingly limited declustering the closer the MM is to the duplex end (Figure 2b). Possible reasons for the limitation in declustering are: 1) $TH_F$ does not detach from Sx due to a) T not reaching the last bp of BM (7), b) T reaching the last bp of BM, but $TH_F$ being too stable to detach spontaneously, potentially initiating a backwards TMSD reaction; or 2) $TH_F$ does dissociate, but rehybridizes to Sx within a short timeframe, leading to no effective change in $V_H$ and no measurable magnetic signal change. To investigate the influence of the second aspect, we added 10 nt blocker strands (table S4) to the assay, which are designed to hybridize to accessible $L_{DSD}$ or Sx strands, blocking domains $TH_F$ (B*) or $TH_F$* (B) and interfere with the reattachment. We then monitored changes in $\chi''$ over 60 min (Figure S4).

**Table S4. Information on blocker strands B and B*.**

| Strand | Sequence 5′ -> 3′ | Complementary to |
|---|---|---|
| B | TCC TGC TAA C | S0 |
| B* | GTT AGC AGG A | $L_{DSD}$ |

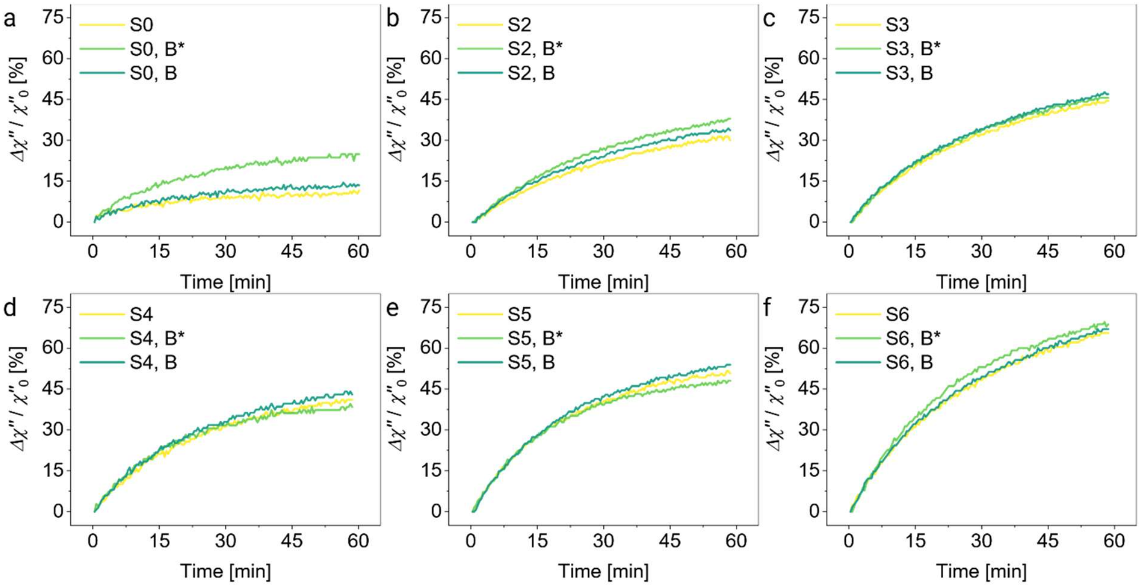

**Figure S4. Declustering in the presence of a blocker strand.** Measured $\Delta\chi''/\chi''_0$ with 100 mM $Mg^{2+}$, 1 nM T (α = 7 bp, β = 6 bp) for all mismatch positions Sx (4 nM) over 60 min in the absence (yellow) and presence of 12 nM blocker strands, offering a ratio of 1:1 for blocker to $L_{DSD}$.

The presence of B* strands introduces a significant signal increase at 60 min for the complementary substrate S0 (panel S4a). This is caused by enhanced declustering, indicating that the B* strands attached to accessible $L_{DSD}$ strands and limited their hybridization to $TH_F$* once S0 and $L_{DSD}$ have separated. These results suggest that for the fully complementary substrate, backwards TMSD may occur. Some minor increase in $\Delta\chi''/\chi''_0$ can be observed for the addition of blocker strands to the clusters with S2 (panel S4b). For the other samples, the presence of blocker strands shows no significant effect, indicating that the limiting factor for the detachment of these DNA duplexes lies within the initial, spontaneous dissociation of $TH_F$.

The trends in signal change stays the same between the mismatch positions Sx in the presence and absence of blocker strands, with S6 showing most signal increase (panel S4f).

#### MATE-based declustering is accelerated by $Mg^{2+}$

In non-amplifying declustering assays prior to the MATE cascade, we observed declustering to be accelerated in the presence of $Mg^{2+}$ (Figure S5a). This is due to the DNA duplex stabilization and enhanced TH formation in the presence of $Mg^{2+}$ ions [19,20]. While we would like to enhance the binding of T to S at $TH_T$, we require $TH_F$ of $L_{DSD}$ to be sufficiently unstable to detach from S once T has completed the BM process. Therefore, we studied the declustering behavior of our MATE-cascade as a function of $Mg^{2+}$ concentrations up to 100 mM after 5 h of incubation without spontaneous dissociation (S0, β = 0 bp, Figure S5b) and over 80 min with spontaneous dissociation in the presence of a mismatch (S6, β = 6 bp, Figure S5c).

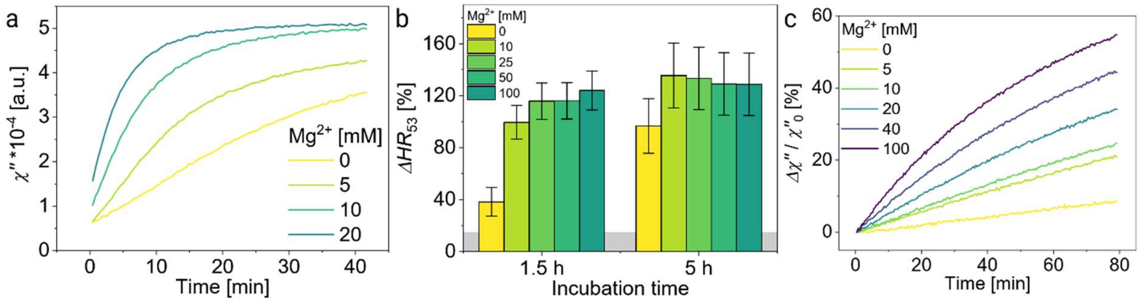

**Figure S5. Acceleration of declustering in the presence of $Mg^{2+}$.** (a) Preliminary $\chi''$ measurement with full displacement of $L_{DSD}$, sequences adapted from [15], measured at 100 Hz, for 4 nM T (α = 7 bp), 3 nM S. (b) $\Delta HR_{53}$ after 1.5 h and 5 h of incubation with 3 nM T (α = 7 bp, β = 0 bp) on 3 nM S0, normalized to a control sample without $Mg^{2+}$ and T. Grey area marks 1σ of control sample. (c) $\Delta\chi''/\chi''_0$ measured on clusters with 2 nM S6 with 1 nM T (α = 7 bp, β = 6 bp) at 120 Hz.

The measured $\Delta HR_{53}$ (Figure S5b) show that all samples containing $Mg^{2+}$ have almost reached their saturation value after 1.5 h (~ 130 %), while the sample without $Mg^{2+}$ significantly lag behind with only ~ 40 % after 1.5 h and ~ 95 % after 5 h.

Monitored over 80 min, the effect of $Mg^{2+}$ can be observed better (Figure S5c). The value of 0 mM $Mg^{2+}$ after 80 min ($\Delta\chi''/\chi''_0$ = 8.7 %) was reached after only 8 min for the sample containing 100 mM $Mg^{2+}$. The measured signal change $\Delta\chi''/\chi''_0$ after 10 min improved by factor 16.9 from 0.7 % for 0 mM to 11.8 % for 100 mM $Mg^{2+}$. We, therefore, argue that the presence of $Mg^{2+}$ improves the hybridization of $TH_T$ more significantly than it inhibits the spontaneous dissociation of $TH_F$.

#### Increasing length of $TH_T$ enhances MATE-based declustering

The effective declustering rate in classic TMSD depends significantly on the length of the invader-TH. The displacement rate typically saturates for TH lengths between 6 to 7 nt [15,21]. We investigated the impact of the $TH_T$ length α on the declustering with the spontaneous dissociation of β = 6 bp, in the absence and presence of 100 mM $Mg^{2+}$ (Figure S6).

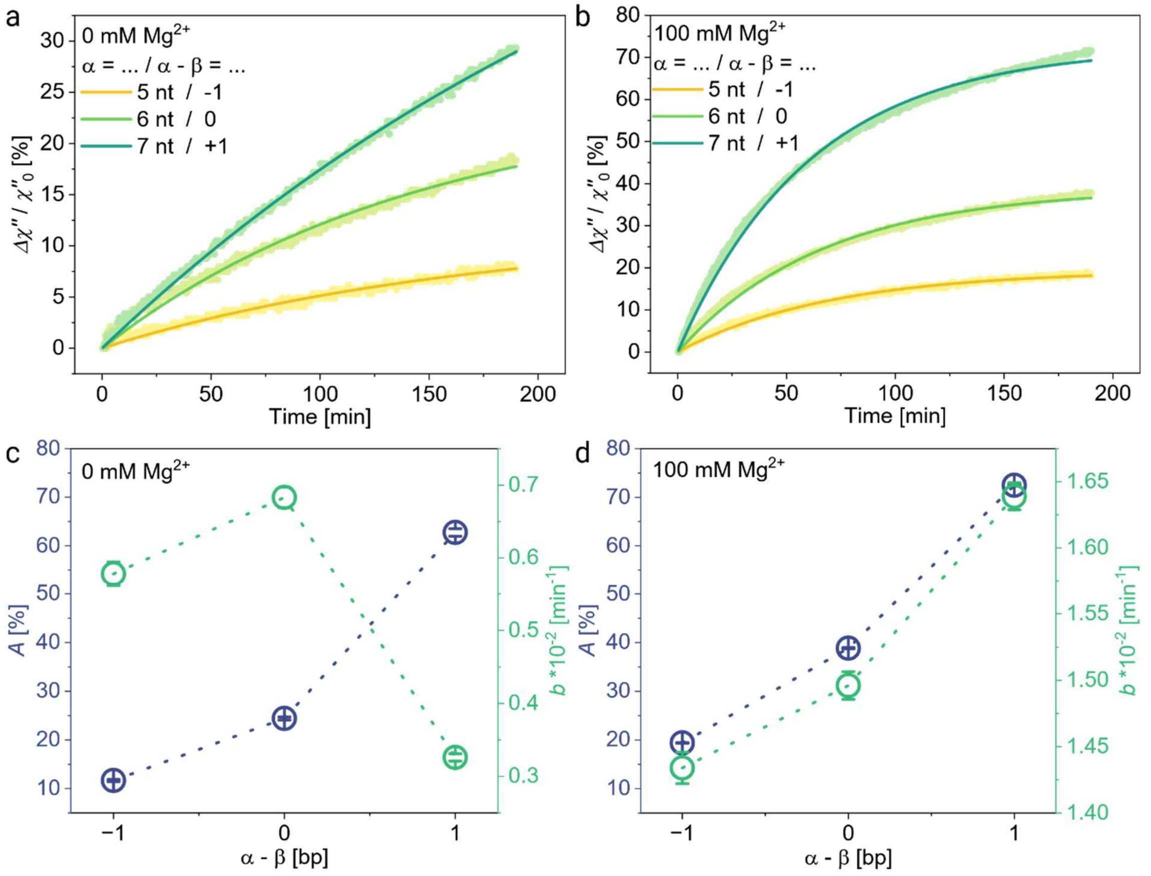

**Figure S6. Influence of $TH_T$ length α on declustering kinetics.** $\Delta\chi''/\chi''_0$ measured on S6 (3 nM), 1 nM T (β = 6 bp), for a) 0 mM and b) 100 mM $Mg^{2+}$. Fit parameters from equation 1 for c) 0 mM $Mg^{2+}$ and d) 100 mM $Mg^{2+}$.

To understand these data sets better, we fitted them to equation 1 and obtained the apparent reaction rate $b$ and $A$ parameters (Figure S6c, d), both determining the sensitivity of the MATE cascade. Rate $b$ increases significantly by adding $Mg^{2+}$ to the assay and is the highest for α = 7 bp at 100 mM $Mg^{2+}$. The parameter $A$ increases by 5.4-fold (0 mM Mg2+) and 3.7-fold (100 mM Mg2+) by changing (α − β) from -1 to 1 (Figure S6c, d). This can be explained by considering the thermodynamics of the system. For the case of α−β = −1 bp, the system loses one bp by declustering (stage 1 to stage 2 (Figure 1)), which is thermodynamically unfavorable. In contrast, by gaining one additional bp in the case of α−β = +1 bp, the declustering processes are thermodynamically favorable, thus resulting in more declustering and signal gain. The effect of $Mg^{2+}$ concentration on the assay sensitivity can be better seen by looking at the maximum signal gain after a certain incubation time (e.g. 2 h, as taken in MPS-based assays) in the absence and presence of $Mg^{2+}$ (Figure S6a, b). In case of α = 7 bp, the signal increases from 11.3 % to 22.8 % after 1 h and from 20.1 % to 31.7 % after 2 h by adding 100 mM $Mg^{2+}$.

#### Longer $TH_F$ length improves target recycling

We first investigated the recycling capability of our MATE-cascade with complementary substrate (S0) for two lengths β of $TH_F$ (Figure S7a).

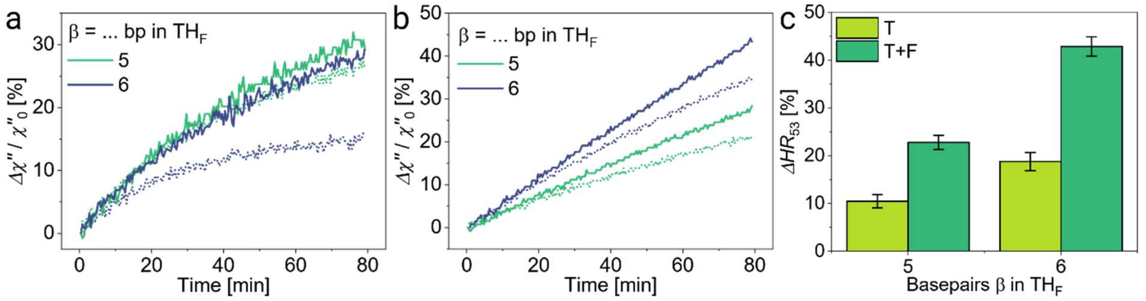

**Figure S7. Recycling is enhanced for longer $TH_F$ domain.** $\Delta\chi''/\chi''_0$ without (dotted) and with 100 nM Fx (solid) for two $TH_F$ lengths β, on clusters with a) complementary substrate (S0) and b) MM at position 5 (S5). c) $\Delta HR_{53}$ after 4 h of incubation with S5, normalized to a control sample with 100 nM F5. Grey area marks 1σ of control sample. (All measurements were executed with 1 nM T (α = 7 bp), 4 nM Sx, 0 and 100 nM Fx, 100 mM $Mg^{2+}$.)

While spontaneous dissociation for the MM-free case S0 is impeded by elongating $TH_F$ from 5 to 6 bp (Figure S7a, dotted lines), the signal increases similarly in the presence of 100 nM F0. For β = 5 bp, $\Delta\chi''/\chi''_0$ at 80 min increases only 3.3 % from 26.3 % (T) to 29.6 % (T+F). For β = 6 bp, the signal change without fuel (T) is only 16.1 %, while the sample with fuel (T+F) shows $\Delta\chi''/\chi''_0$ = 29.3 %. The increased recycling can be observed in the presence of a MM (S5) as well, measured over 80 min (Figure S7b) and after 4 h (Figure S7c). The observed $\Delta HR_{53}$ increases from 10.5 % (T) to 22.8 % (T+F) for β = 5 bp. For β = 6 bp, both values increase almost two-fold to 18.8 % (T) and 42.9 % (T+F), showcasing the importance of a longer $TH_F$ domain for enhanced recycling.

Increasing the length β of $TH_F$ offers a longer, more stable TH to the reverse TMSD reaction. As a first benefit, the fuel DNA is able to attach to the substrate more strongly and initiate the recycling of the incumbent target. The second advantage lies in the nature of the TH exchange, where upon dissociation of the 7 bp of $TH_T$, 2 bp are lost for β = 5 bp, while the duplex is deprived of only 1 bp for β = 6 bp. These two factors combined enhance the reverse TMSD reaction and the related recycling of the target DNA.

#### High fuel concentrations are required for fast target recycling

Increasing concentrations of fuel were added to clusters (4 nM S6, 1 nM T (α = 7 bp, β = 6 bp), 100 mM $Mg^{2+}$) and monitored over 80 min (Figure S8), showing the importance of oversaturated fuel concentrations for effective recycling to take place.

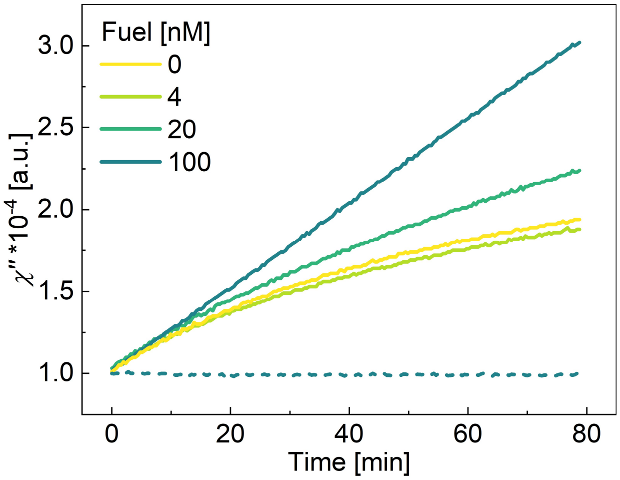

**Figure S8. Concentration driven recycling enhances MATE efficiency.** $\chi''$ was measured in the presence (solid lines) and absence (dashed line) of 1 nM T.

#### Leakage declustering depends on the mismatch position x

Leakage-based declustering was monitored for all MM positions Sx (4 nM) for two additional fuel concentrations to Figure 3c with 100 mM $Mg^{2+}$ (Figure S9), showing the same trend in leakage over all three sets of measurements.

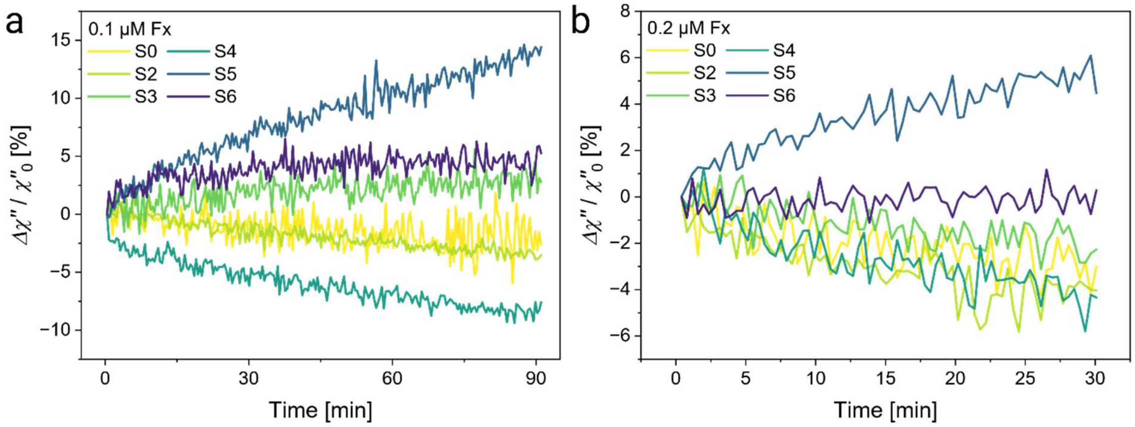

**Figure S9. Leakage-based declustering for all mismatch positions x.** a) 0.1 µM Fx, b) 0.2 µM Fx.

#### NUPACK simulations of DNA complexes

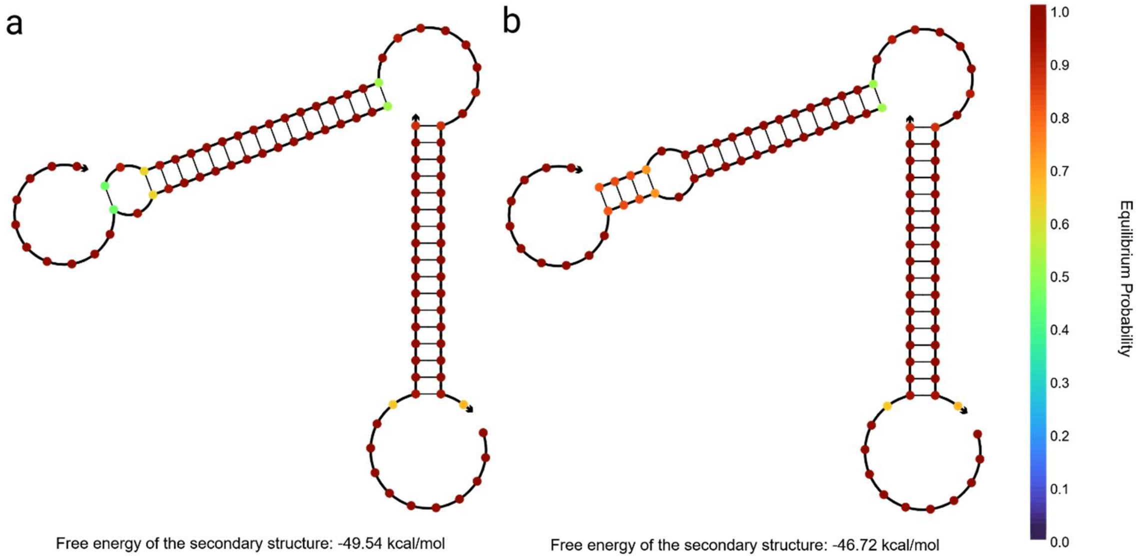

**Figure S10. NUPACK simulations of DNA complexes $L_{Stay}$ + $L_{DSD}$ + Sx.** Simulation parameters: 150 mM $Na^{+}$, 0 mM $Mg^{2+}$, 5 nM Oligo, 25 °C, all stacking. a) S2, b) S5.

#### Sequences for CG-variation

**Table S5. DNA sequences used for CG-variation in Figure 4.** These sequences replace the respective DNA strands during cluster preparation and measurement.

| Name | Sequence (5´ -> 3´) |
|---|---|
| L_DSD,GC | ACC CGC AAT CCT GCT ACG (AAA AAA AAA A) [BIOTEG] |
| S_6,GC | CGT AGA AGG ATT GCG GGT GCC AAT GAG CTA CAC ACA TAC AAT TA |
| F_6,GC | ACC CGC AAT CCT TCT ACG |

#### Zoom on first 60 minutes for target series

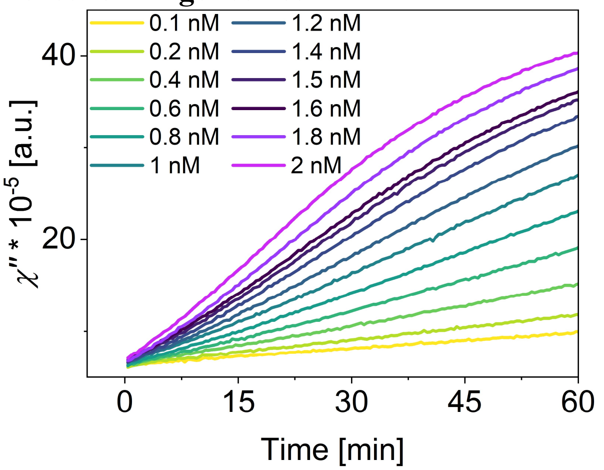

**Figure S11. Signal lag for increasing target concentrations.** Unprocessed $\chi''$ from Figure 5b, zoomed on the first 60 min. 4 µM F, 100 mM $Mg^{2+}$, 4 nM S6 LNA7A.

### References

(1) Mohammadniaei, M.; Zhang, M.; Ashley, J.; Christensen, U. B.; Friis-Hansen, L. J.; Gregersen, R.; Lisby, J. G.; Benfield, T. L.; Nielsen, F. E.; Henning Rasmussen, J.; Pedersen, E. B.; Olinger, A. C. R.; Kolding, L. T.; Naseri, M.; Zheng, T.; Wang, W.; Gorodkin, J.; Sun, Y. A Non-Enzymatic, Isothermal Strand Displacement and Amplification Assay for Rapid Detection of SARS-CoV-2 RNA. *Nat. Commun.* **2021**, *12* (1). https://doi.org/10.1038/s41467-021-25387-9.

(2) SantaLucia, J.; Hicks, D. The Thermodynamics of DNA Structural Motifs. *Annu. Rev. Biophys. Biomol. Struct.* **2004**, *33*, 415–440. https://doi.org/10.1146/annurev.biophys.32.110601.141800.

(3) Abraham Punnoose, J.; Thomas, K. J.; Chandrasekaran, A. R.; Vilcapoma, J.; Hayden, A.; Kilpatrick, K.; Vangaveti, S.; Chen, A.; Banco, T.; Halvorsen, K. High-Throughput Single-Molecule Quantification of Individual Base Stacking Energies in Nucleic Acids. *Nat. Commun.* **2023**, *14* (1). https://doi.org/10.1038/s41467-023-36373-8.

(4) Chowdhury, M. S.; Rösch, E. L.; Esteban, D. A.; Janssen, K. J.; Wolgast, F.; Ludwig, F.; Schilling, M.; Bals, S.; Viereck, T.; Lak, A. Decoupling the Characteristics of Magnetic Nanoparticles for Ultrahigh Sensitivity. *Nano Lett.* **2023**, *23* (1), 58–65. https://doi.org/10.1021/acs.nanolett.2c03568.

(5) Chugh, V. K.; di Girolamo, A.; Krishna, V. D.; Wu, K.; Cheeran, M. C. J.; Wang, J. P. Frequency and Amplitude Optimizations for Magnetic Particle Spectroscopy Applications. *J. Phys. Chem. C* **2023**, *127* (1), 450–460. https://doi.org/10.1021/acs.jpcc.2c07534.

(6) Wolgast, F. T.; Kahmann, T.; Janssen, K. J.; Yoshida, T.; Zhong, J.; Schilling, M.; Ludwig, F.; Viereck, T. Low-Cost Magnetic Particle Spectroscopy Hardware for Low-Viral-Load Immunoassays. *IEEE Trans. Instrum. Meas.* **2024**. https://doi.org/10.1109/TIM.2024.3449985.

(7) Ludwig, F.; Guillaume, A.; Schilling, M.; Frickel, N.; Schmidt, A. M. Determination of Core and Hydrodynamic Size Distributions of CoFe 2O4 Nanoparticle Suspensions Using Ac Susceptibility Measurements. *J. Appl. Phys.* **2010**, *108* (3). https://doi.org/10.1063/1.3463350.

(8) Narayanasamy, K. K.; Cruz-Acuña, M.; Rinaldi, C.; Everett, J.; Dobson, J.; Telling, N. D. Alternating Current (AC) Susceptibility as a Particle-Focused Probe of Coating and Clustering Behaviour in Magnetic Nanoparticle Suspensions. *J. Colloid Interface Sci.* **2018**, *532*, 536–545. https://doi.org/10.1016/j.jcis.2018.08.014.

(9) Rösch, E. L.; Zhong, J.; Lak, A.; Liu, Z.; Etzkorn, M.; Schilling, M.; Ludwig, F.; Viereck, T.; Lalkens, B. Point-of-Need Detection of Pathogen-Specific Nucleic Acid Targets Using Magnetic Particle Spectroscopy. *Biosens. Bioelectron.* **2021**, *192*. https://doi.org/10.1016/j.bios.2021.113536.

(10) Rösch, E. L.; Sack, R.; Chowdhury, M. S.; Wolgast, F.; Zaborski, M.; Ludwig, F.; Schilling, M.; Viereck, T.; Rand, U.; Lak, A. Amplification- and Enzyme-Free Magnetic Diagnostics Circuit for Whole-Genome Detection of SARS-CoV-2 RNA. *ChemBioChem* **2024**, *25* (16). https://doi.org/10.1002/cbic.202400251.

(11) Allen, R. J.; Valeriani, C.; Rein Ten Wolde, P. Forward Flux Sampling for Rare Event Simulations. *J. Phys. Condens. Matter* **2009**, *21* (46). https://doi.org/10.1088/0953-8984/21/46/463102.

(12) Snodin, B. E. K.; Randisi, F.; Mosayebi, M.; Šulc, P.; Schreck, J. S.; Romano, F.; Ouldridge, T. E.; Tsukanov, R.; Nir, E.; Louis, A. A.; Doye, J. P. K. Introducing Improved Structural Properties and Salt Dependence into a Coarse-Grained Model of DNA. *J. Chem. Phys.* **2015**, *142* (23). https://doi.org/10.1063/1.4921957.

(13) Poppleton, E.; Matthies, M.; Mandal, D.; Romano, F.; Šulc, P.; Rovigatti, L. OxDNA: Coarse-Grained Simulations of Nucleic Acids Made Simple. *J. Open Source Softw.* **2023**, *8* (81), 4693.

https://doi.org/10.21105/joss.04693.

(14) Ouldridge, T. E.; Šulc, P.; Romano, F.; Doye, J. P. K.; Louis, A. A. DNA Hybridization Kinetics: Zippering, Internal Displacement and Sequence Dependence. *Nucleic Acids Res.* **2013**, *41* (19), 8886–8895. https://doi.org/10.1093/nar/gkt687.

(15) Srinivas, N.; Ouldridge, T. E.; Šulc, P.; Schaeffer, J. M.; Yurke, B.; Louis, A. A.; Doye, J. P. K.; Winfree, E. On the Biophysics and Kinetics of Toehold-Mediated DNA Strand Displacement. *Nucleic Acids Res.* **2013**, *41* (22), 10641–10658. https://doi.org/10.1093/nar/gkt801.

(16) Sengar, A.; Ouldridge, T. E.; Henrich, O.; Rovigatti, L.; Šulc, P. A Primer on the OxDNA Model of DNA: When to Use It, How to Simulate It and How to Interpret the Results. *Front. Mol. Biosci.* **2021**, *8* (June), 1–22. https://doi.org/10.3389/fmolb.2021.693710.

(17) Sample, M.; Liu, H.; Matthies, M.; Šulc, P. Hairygami: Analysis of DNA Nanostructures' Conformational Change Driven by Functionalizable Overhangs. **2023**. https://doi.org/10.1021/acsnano.4c10796.

(18) Cisse, I. I.; Kim, H.; Ha, T. A Rule of Seven in Watson-Crick Base-Pairing of Mismatched Sequences. *Nat. Struct. Mol. Biol.* **2012**, *19* (6), 623–627. https://doi.org/10.1038/nsmb.2294.

(19) Every, A. E.; Russu, I. M. Influence of Magnesium Ions on Spontaneous Opening of DNA Base Pairs (Journal of Physical Chemistry B (2008) 112B). *J. Phys. Chem. B* **2008**, *112* (47), 15261. https://doi.org/10.1021/jp8091732.

(20) Owczarzy, R.; Moreira, B. G.; You, Y.; Behlke, M. A.; Wälder, J. A. Predicting Stability of DNA Duplexes in Solutions Containing Magnesium and Monovalent Cations. *Biochemistry* **2008**, *47* (19), 5336–5353. https://doi.org/10.1021/bi702363u.

(21) Simmel, F. C.; Yurke, B.; Singh, H. R. Principles and Applications of Nucleic Acid Strand Displacement Reactions. *Chem. Rev.* **2019**, *119* (10), 6326–6369. https://doi.org/10.1021/acs.chemrev.8b00580.
